# Assembly and dynamics of the outer membrane exopolysaccharide transporter PelBC of *Pseudomonas aeruginosa*

**DOI:** 10.1101/2024.09.23.614597

**Authors:** Marius Benedens, Cristian Rosales Hernandez, Sabine A.P. Straathof, Jennifer Loschwitz, Otto Berninghausen, Giovanni Maglia, Roland Beckmann, Alexej Kedrov

## Abstract

The infamous opportunistic pathogen *Pseudomonas aeruginosa* enhances its virulence and antibiotic resistance upon formation of durable biofilms. The biofilm stability is mediated by its matrix built of secreted exopolysaccharides, eDNA, and structural proteins. Exopolysaccharides of *P. aeruginosa* – Pel, Psl and alginate – have the highest biomedical relevance, but the mechanisms behind their synthesis and secretion are poorly understood. Here, we employ cryogenic electron microscopy to resolve the 2.5 Å structure of the outer membrane complex PelBC for Pel exopolysaccharide, which is uniquely composed of the membrane-embedded β-barrel PelB and the asymmetrical ring of 12 lipoproteins PelC at the periplasmic interface. The assembly captured in the lipid-based nanodisc is stabilized by electrostatic contacts of PelC with the periplasmic loops of PelB and multiple interactions with PelB N-terminal helical domains. Within the membrane, the resolved acyl chains of the PelC lipoproteins are alternated by the tryptophan residues immersed into the lipid leaflet, thus offering a stable anchoring architecture. The highly anionic interior of the PelB β-barrel is sealed by three loops at the extracellular side, where the short Plug-S loop is aligned with the periplasmic helical scaffold, being the potential gating element for the Pel exopolysaccharide tunneling. Molecular dynamic simulations of PelB in native-like membrane environments suggest that Plug-S is sufficiently flexible to open a tunnel, and so serve as a gate. The gating model is further supported by single-channel conductivity measurements, which identify two conductance states of PelB. Via mutational analysis we confirm that Plug-S mediates opening of a narrow tunnel, as required for the controlled exopolysaccharide transport. Our structural and functional analysis of the pathogenicity-relevant complex offer a detailed and comprehensive view on this unique machinery and suggest the route taken by the exopolysaccharide at the final secretion step.

## Introduction

The Gram-negative bacterium *Pseudomonas aeruginosa* is an opportunistic human pathogen that accounts for nearly 20 % of all nosocomial infections, being a major risk factor for immunocompromised patients and those with cystic fibrosis (Nelson *et al*, 2021). *P. aeruginosa* infections build up a burden for the healthcare systems worldwide due to the prolonged hospitalization period and the associated costs. The primary challenges to combat this versatile bacterium are its rapidly developing antibiotic resistance and formation of extensive durable biofilms, both in tissues upon host invasion and on diverse abiotic surfaces. Stability of the biofilms is mediated by the complex composition of the matrix, where the exopolysaccharides - alginate, Psl and Pel - serve to embed cells, ensure the mechanical strength of the matrix and provide attachment sites for secreted virulence factors (Gheorghita *et al*, 2023). Despite the highest biomedical relevance of the exopolysaccharides of *P. aeruginosa*, the understanding of the molecular mechanisms of their secretion is poor.

Pel is the major constituent of *P. aeruginosa* pellicles, i.e. films formed at the water-air interface, and it plays a key role at early stages of the biofilm formation at solid surfaces. Pel has been recently identified in other b-, d-, and g-proteobacteria, but also several extremophiles and Gram-positive bacteria (Whitfield *et al*, 2020a). *P. aeruginosa* Pel is composed of 1-4 a-linked N-acetyl-D-galactosamine residues, which are partially deacetylated in the periplasm, so the Pel exopolysaccharide acquires positive charge along the secretion pathway (Le Mauff *et al*, 2022). This charge is utilized then to crosslink the extracellular DNA thus contributing to the matrix assembly (Jennings *et al*, 2015). Other roles of Pel involve stabilization of the adhesin CdrA and complementation for defects in the outer membrane, as deletion of the major structural protein OprF stimulates Pel accumulation at the cell surface (reviewed in (Gheorghita *et al*., 2023)).

Secretion of Pel in the Gram-negative *P. aeruginosa* assumes the polysaccharide translocation across the inner and outer membranes (Figure 1), and the genes essential for the process are encoded in a single *pelABCDEFG* operon. The polysaccharide elongation mediated by the glycosyltransferase PelF in the cytoplasm is coupled to the synthase-dependent translocation through the inner membrane complex of PelD, PelE and PelG subunits, in a still unknown fashion (Figure 1) (Whitfield *et al*, 2020b). Partial deacetylation of Pel in the periplasm is mediated by PelA, prior crossing the outer membrane through PelB (Marmont *et al*, 2017b). PelB is composed of a C-terminal β-barrel preceded by the helical scaffold, built of multiple tetratricopeptide repeats (TPR). The length of the modelled scaffold exceeds 20 nm, thus being sufficiently long to span the periplasm (Suppl. Figure 1A). Notably, the N-terminal end of PelB contains 22 apolar and aromatic residues, which may form a transmembrane helix within the inner membrane without a cleavage site for the signal peptidase (Suppl. Figure 1B). Thus, PelB of *P. aeruginosa*, as well as its multiple homologs in other species, may physically bridge two membranes, with the TPR scaffold forming a passage for the polysaccharide across the periplasm.

**Figure 1.**
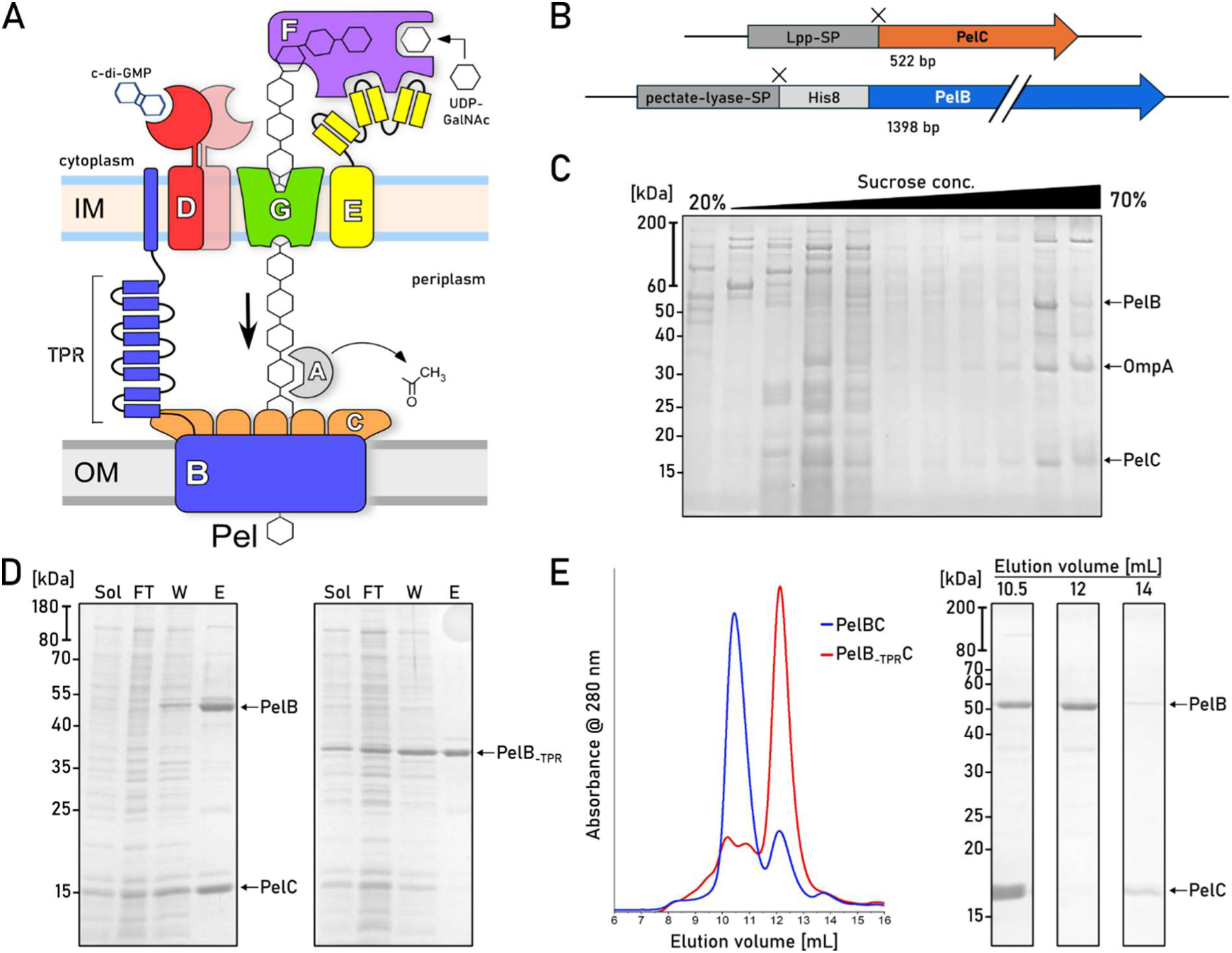
Isolation of the natively assembled PelBC complex of *P. aeruginosa*. (A) Putative organization of the synthesis/secretion machinery of the Pel exopolysaccharide in *P. aeruginosa*. (B) Gene constructs of PelB and PelC in the pETDuet-1 vector. Cleavage sites after the signal peptides of both proteins are indicated as “X”. (C) SDS-PAGE of the membrane isolates after the sucrose gradient shows PelB and PelC co-occurring in late fractions characteristic for the outer membrane vesicles. A reference band for the outer membrane protein A (OmpA) is indicated. (D) SDS-PAGE of immobilized metal affinity chromatography on PelBC with His-tagged PelB variants, wild-type (left) and the truncated barrel lacking the periplasmic helical scaffold (right, PelB_-TPR_). Loaded fractions: “Sol.” -detergent-solubilized material; “FT” -flow-through; “W” - wash, “E” - elution. (E) Left: Size-exclusion chromatograms of the PelBC samples from (D). Right: SDS-PAGE of the major SEC fractions collected for the wild-type PelBC.

The β-barrel of PelB is assumed to form a complex with PelC lipoproteins, a hallmark of the Pel system in Gram-negative bacteria. A partial crystal structure of PelC from *Paraburkholderia phytofirmans* solved in the absence of the membrane and the lipid anchors revealed the protomers arranged into a dodecameric ring with a 3.2 nm-wide pore in the center (Marmont *et al*, 2017a). The dimension of the PelC pore matches closely the diameter of the modelled PelB structure (Suppl. Figure 2), and the negative charge on the periplasm-facing side of PelC was suggested to assist in translocation of the cationic polysaccharide. In absence of the complete PelBC structure, AlphaFold-based modelling of the complex in 1:12 stoichiometry provides a high-confidence assembly, where the TPR domain is placed within the central pore of PelC ring. However, models of a similar confidence could also be rendered at different stoichiometries, either having less PelC subunits (ratio 1:11) or more (ratios 1:13 and above) (Suppl. Figure 2). Since the assemblies of lipoproteins in solution and at the membrane interface may differ (Goyal *et al*, 2014), visualizing the actual architecture of the PelBC complex remains an open task. Several mutations within PelC were identified which inhibited the biofilm formation, either via direct interactions with the polysaccharide or their involvement in the PelBC complex assembly (Marmont *et al*., 2017a). Here, the experimentally determined structure should explain their role, and potentially address protein:lipid interactions for the lipoproteins and the transmembrane β-barrel. Critically, the modelled PelB barrel does not manifest a substantially wide conduit at the extracellular side for the polysaccharide translocation, being sealed by several loops. Though the model may reflect the idle state of the channel, the experimental structure determination is required to understand the functional dynamics of the complex.

Here, we employed cryogenic electron microscopy (cryo-EM) to visualize the structure of the PelBC complex of *P. aeruginosa* in the lipid-based nanodiscs, reaching a final resolution of 2.5 Å. Next to the overall architecture of the asymmetric complex of 250 kDa, we report extensive protein:lipid interactions formed by both the lipoproteins and the β-barrel and discover that the periplasm-exposed helical domains of PelB and the highly conserved tryptophan of PelC are essential for the assembly of the complex. The acquired structure suggests the route of the exopolysaccharide across the channel, and it is further used to design single-channel conductivity experiments and computational modelling, which jointly provide first evidence for PelB conformational dynamics in the lipid membrane.

## Results

### Isolation and nanodisc reconstitution of the intact PelBC complex

To establish heterologous expression of the PelBC complex, both genes were cloned into pETDuet-1 vector under individual T7 promotors. To facilitate the efficient export of the synthesized proteins into the periplasm via the Sec machinery, the signal sequence of PelC was substituted with the signal sequence of Lpp, the highly abundant Braun’s lipoprotein of *E. coli*. PelB was expressed as the C-terminal fragment (residues 762-1193) containing three periplasmic TPR repeats and the transmembrane β-barrel, which were preceded with the conventional signal sequence of pectate lyase B of *Erwinia carotovora* and the octa-histidine tag (Figure 1B). Once expressed in *E. coli* C41(DE3) *ΔompF ΔacrAB* strain (Kanonenberg *et al*, 2019), the protein localization to the outer membrane was validated by the ultracentrifugation in the sucrose density gradient: Both proteins appeared within the high-density fraction together with OmpA, an intrinsic marker for the outer membrane vesicles (Figure 1C) (Kanonenberg *et al*., 2019).

Both PelB and PelC were extracted from the membranes with the mild non-ionic detergent DDM. IMAC based on the N-terminal histidine-tag of PelB resulted in co-purification of substantial amounts of PelC (Figure 1D), suggesting that the proteins resided as a complex. The PelB-PelC isolates manifested three distinct peaks in size exclusion chromatography (SEC, Figure 1E): The major peak contained both PelB and the excess of PelC and it was observed at 10.5 mL elution volume suggesting the molecular weight of ∼300 kDa, while the downstream peaks at 12 mL and 14 mL contained PelB and PelC, respectively. The appearance of the tag-less free PelC subunits in the latter peak indicated partial disassembly of the detergent-solubilized complex after the IMAC stage. To prevent that, SEC-purified PelBC complexes were immediately reconstituted into nanodiscs in presence of POPC:POPG lipids (molar ratio 70:30). The chosen scaffold protein MSP1D1 builds nanodiscs of ∼8 nm inner diameter (Hagn *et al*, 2013) that should be sufficient to accommodate the β-barrel of PelB and the acyl chain anchors of PelC lipoproteins, assuming the ring-shaped assembly. SEC of PelBC-nanodiscs resulted in a major peak at ∼10.5 mL where PelB, PelC and MSP1D1 were found, followed by minor amounts of PelB-only nanodiscs (Figure 2A). The lipid environment of nanodiscs greatly stabilized the complex, as no dissociation of PelC was observed in SEC, so the lipoprotein remained anchored within the lipid layer.

**Figure 2.**
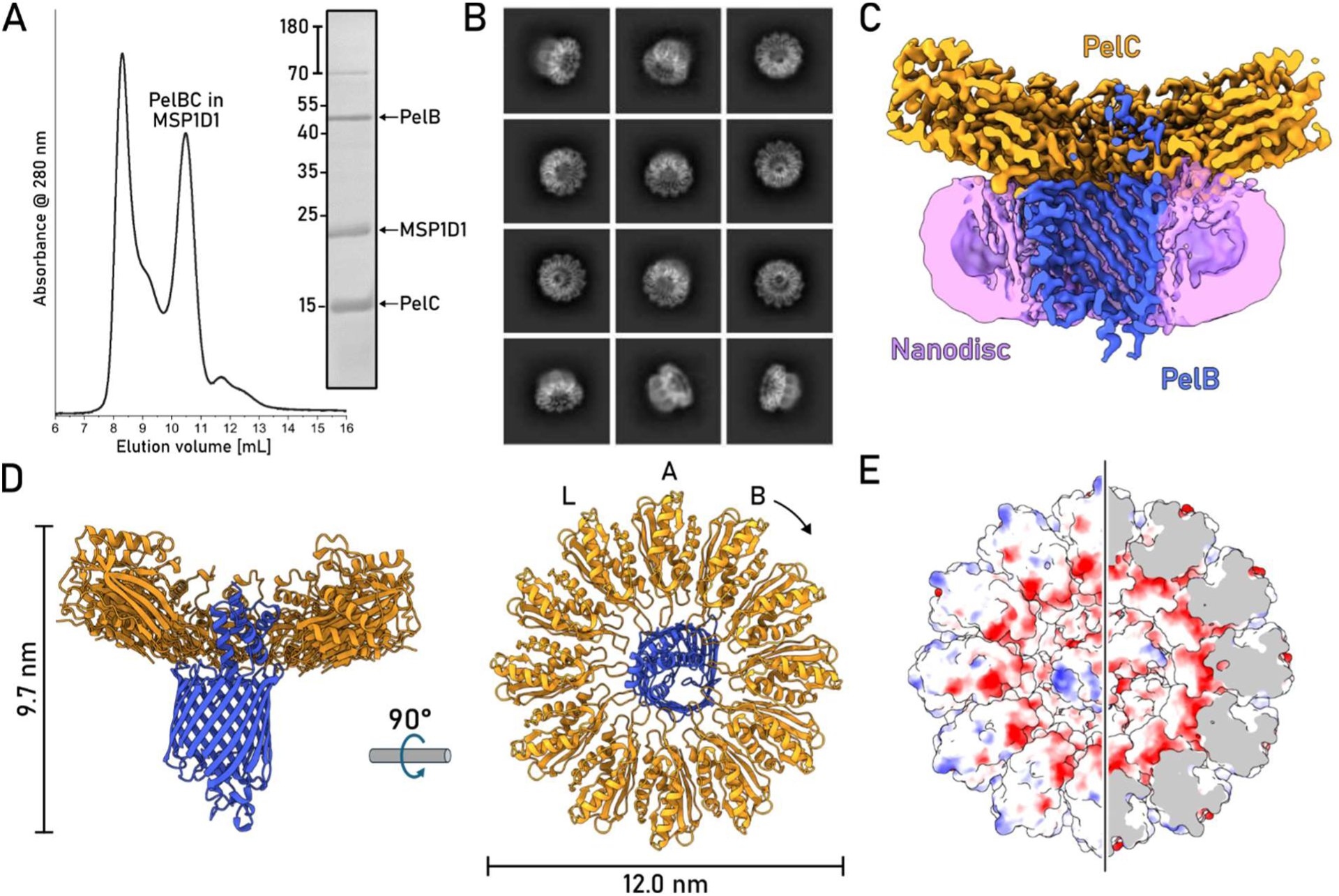
Structure of the nanodisc-embedded PelBC. (A) SEC of the PelBC nanodisc preparation and SDS-PAGE of the fraction at 10.5 mL elution volume used for cryo-EM analysis. (B) Characteristic 2D classes found upon cryo-EM data processing show various projections of the assembled PelBC complexes in the nanodiscs. (C) Final 3D reconstruction (cross-section) of the PelBC complex in the nanodisc with structural elements colored. The (D) The structural model of PelBC complex. PelC subunits labeled from “A” to “L”, so the subunit A approaches the N-terminal end of the PelB β-barrel. The Plug-S loop occupying the exit tunnel of PelB is seen in the periplasmic view (right). (E) The electrostatic potential at the periplasmic surface of the PelC ring (left) and within the inner cavity under the capping loop (right).

### Cryo-EM resolves the architecture of the PelBC complex in the lipid bilayer

The nanodisc-stabilized PelBC complexes were subjected to cryogenic electron microscopy (cryo-EM). From around 30,000 movies, 4.8 million particles were picked for downstream processing. In reference-free 2D class averages, a balanced distribution of side and top views of the PelBC complex was observed (Figure 2B and Suppl. Figure 3), and the nanodisc density could be well-recognized in side views, with the outer diameter of 9 nm in agreement with the previous reports (Ritchie *et al*, 2009). Notably, no classes displaying stoichiometry different from 1:12 PelB:PelC were obtained. The homogeneity of the particle distribution allowed for a 3D reconstruction of the PelBC complex at 2.5 Å resolution with no symmetry imposed (Suppl. Figure 3 and 4). Extensive sorting was carried out to look for different conformations of the complex, but no major structural differences between the classes were detected. A focused classification on PelB resulted in a class with the best resolved β-barrel with short α-helical loops and the N-terminal helical extension, which subsequently led to the final map used for model building (Figure 2C and D). In agreement with the prediction, the complex is composed of a single PelB barrel built of 16 antiparallel β-strands embedded in the lipid bilayer and a dodecameric ring of PelC lipoproteins mounted at the periplasmic side (Figure 2E). Two TPR domains resolved from the residues Gly-803 and the intermediate helix (“stalk”) of PelB are located within the ring, being docked at multiple PelC subunits, as described below. The residues ranging from Asn-762 to Ile-802 remained unresolved, likely due to their higher flexibility. PelB barrel is centrally positioned within the nanodisc, without contacts to the edges which could otherwise distort the conformation. The architecture of PelB is strikingly similar to bacterial transporters for cellulose (BcsC, (Acheson *et al*, 2019)) and PNAG (PgaA, (Wang *et al*, 2016)), which share the overall fold of the 16-stranded β-barrel, with partially structured loops occluding the extracellular exit of the channel (Suppl. Figure 5).

The interior of the PelB β-barrel is predominantly negatively charged, where the charge is distributed asymmetrically, being largely concentrated on strands 11 to 14 (Figure 3A). The anionic cluster faces the groove within the TPR domains, so the cationic Pel polysaccharide may employ this route for entering the channel driven by the electrostatic interactions. The anionic wall is continued with the negatively charged and partially structured loops forming the bottom of the barrel at the extracellular side with no tunnel sufficiently wide for the EPS passage (Figure 3B and C). As a major element here, a short a-helix between the strands 7 and 8 (further referred as Plug-I) is bent inwards the central cavity of PelB where it lays perpendicular to the barrel axis (Figure 3C). Plug-I forms several contacts within the barrel, first of all via Arg-999 to Tyr-922 in β-strand 2 and Glu-935 in β-strand 3, and so it may serve to stabilize the barrel. Plug-I is opposed by a glycine-rich loop between the C-terminal β-strands 15 and 16 (Plug-S), and together these two loops occlude the exit tunnel of PelB (Figure 3B). The loop between the β-strands 11 and 12 (Plug-O) is exposed to the solvent at the extracellular side forming a “dome” over the barrel. Though only a small part of this 31 residues-long loop builds an α-helix, the polypeptide chain is well-resolved in cryo-EM, so Plug-O is rigid at the experimental conditions. Comparison of the resolved *P. aeruginosa* PelB structure with the models of PelB homologs from other *Pseudomonas* species shows high conservation of the extracellular loops, apart the Plug-O (Suppl. Figure 6 and 7). Here, broad variations in the length and the putative structures are observed, suggesting that the loop architecture evolves in response to the certain habitat of the bacteria and/or its specie-specific LPS layer.

**Figure 3.**
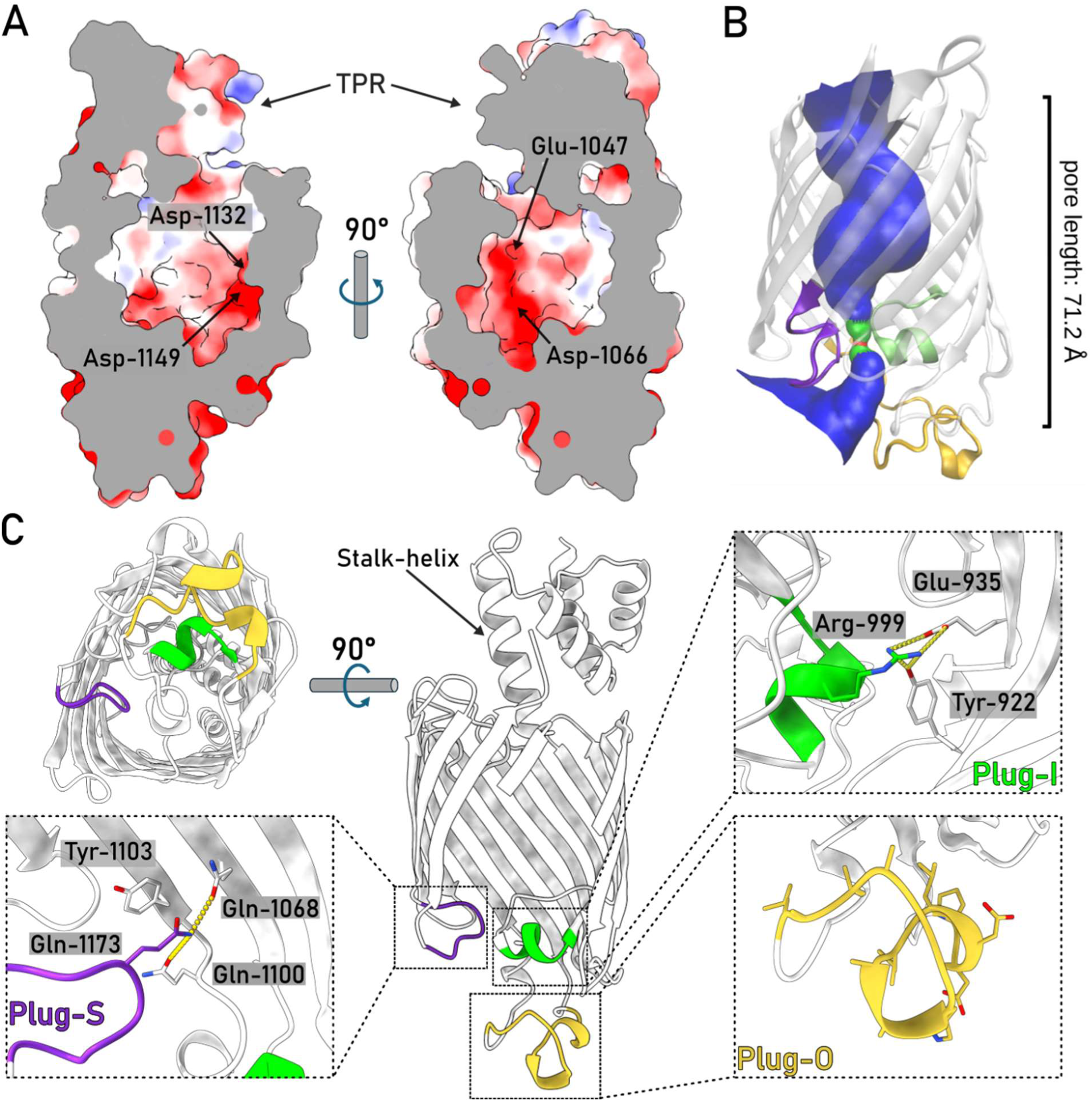
The architecture of the PelB β-barrel. (A) The charge distribution inside the PelB β-barrel is asymmetric, with anionic residues localized at the β-strands 11-14 and the bottom of the barrel. Representative residues are indicated. (B) The loops at the extracellular side of the β-barrel serve to seal the pore. The tunnel is rendered by HOLE algorithm, the bottleneck impermeable for water molecules is indicated in red. (C) Detailed view on the extracellular loops of PelB and the interactions which stabilize their positions in the resolved structure. The universally conserved residue Tyr-1103 near the Plug-S loop is indicated.

Similar to the barrel interiors (Figure 3), the extracellular side of *P. aeruginosa* PelB is highly anionic with the net charge of −11 (Asp+Glu: −18, Arg+Lys: 7), which appears as a common feature among PelB homologs in *Pseudomonas* species (Suppl. Figure 5). Such charge distribution is also seen for several *P. aeruginosa* porins, incl. PA4067 (OprG; PDB ID 2X27), PA3280 (OprO; 4RJX), and PA3279 (OprP, 2O4V), and the negative charge may serve to repel the phosphate groups of the surrounding LPS cores. In context of the polysaccharide secretion, it may also be involved in functional opening/closing of the barrel, and/or providing the electrostatic driving force for translocation of the cationic Pel. For the outer membrane transporters of partially cationic polysaccharides, such as bacterial phosphoethanolamine cellulose and PNAG, the electrostatic interactions at the exit of the secretion machinery were proposed to facilitate the directed transport (Wang *et al*., 2016).

As a unique feature of the Pel translocation system, twelve PelC subunits assemble in a ring at the periplasmic side of the PelB barrel (Figures 2 and 4), and so they repeat the architecture of the crystallized PelC of *P. phytofirmans* (Marmont *et al*., 2017a). Beneficially, the cryo-EM map visualizes the complete periplasm-facing loop-helix turn (residues 70-98) that was not resolved in the crystal structure, due to its flexibility apparent from the current reconstruction (Figure 4A and Suppl. Figure 4). The loop-helix segment caps the PelC subunit beneath and shields the excessive negative charge rendered by the glutamates in positions 52 and 55, but also provides negative charge via Asp-79 and Asp-84 lining the central pore (Figure 4A). At the membrane interface, the extended loop of PelC (further referred as D-loop, for the conserved Asp-119 within) is exposed into the central pore. The electronegative potential rendered by Asp-119 was previously related to PelC:exopolysaccharide interactions (Marmont *et al*., 2017a), as altering the charge here abolished the biofilm formation. However, within the assembled complex those aspartates are buried at the PelB-PelC interface and build multiple contacts with the surface-exposed arginine residues of PelB, such as 905, 944, 1018, 1114, 1122, and 1190 (Figure 4B and C), thus playing rather a structural role. Noteworthy, with glycine residues in positions 117 and 120, the D-loop is sufficiently flexible, so individual PelC subunits adapt their conformations to the proximate structural features of PelB, i.e. the helical domain and the periplasmic loops (Figure 4B).

**Figure 4.**
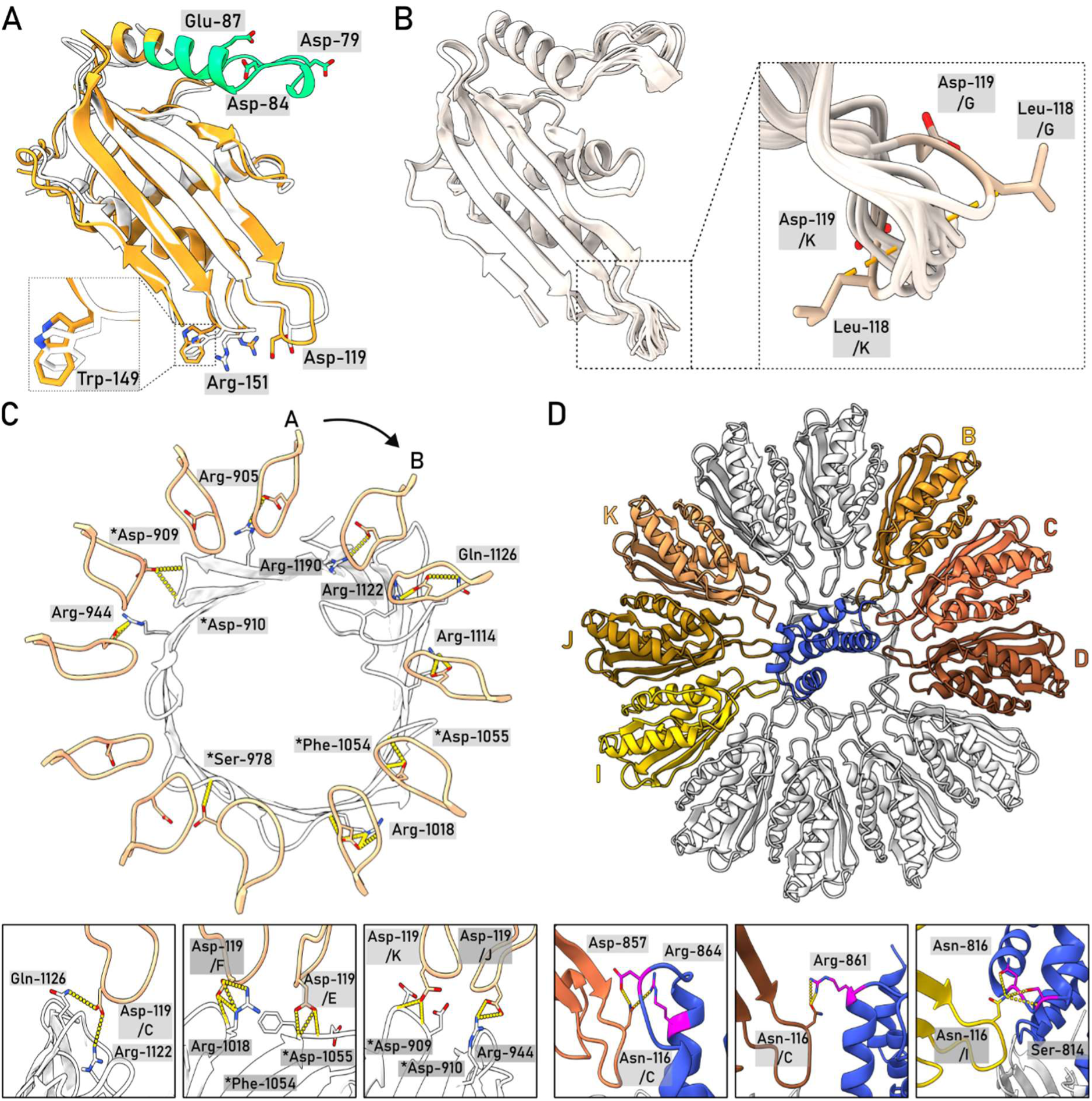
The architecture of the PelC ring and PelC:PelB interactions. (A) Cryo-EM structure of *P. aeruginosa* PelC, subunit E (orange), aligned with the partial crystal structure of PelC from *P. phytofirmans* (white). The capping loop absent in the crystal structure is shown in green. (B) Alignment of 12 PelC subunits resolved in the cryo-EM structure. Due to the subunit-specific interactions with PelB, the pore-exposed D-loops manifest different conformations (zoomed), where the deviations measured for Leu-118 Cβ reach up to 8.5 A for subunits K and G. (C) Asp-119 of PelC subunits is involved in interactions with the periplasmic loops of PelB (residues indicated). PelB residues which interact via the backbone amide group are indicated with asterisk. (D) Multiple PelC subunits (colored) build electrostatic interactions with the helical scaffold of PelB (shown in blue). Subunits C and D interact with the stalk helix, and subunits B and I-K with the TPR domain.

Two PelC subunits, B and C, build electrostatic interactions with the stalk helix of PelB (Figure 4D), and the preceding TPRs are stabilized in defined positions by interactions with PelC subunits B/C/-D and I/J/-K, so these two repeats are well-resolved within the central pore. To test the role of the helical region in the PelBC assembly and stability, we co-expressed PelC with the truncated PelB variant containing only the β-barrel domain. Though both proteins could be extracted from the membrane, no PelC was co-purified with PelB in absence of the stalk helix and the TPR domains (Figure 1E), suggesting that the conserved domain is crucial for the protein:protein interaction, and it may serve as a nucleation site for the PelC ring assembly.

### The PelBC complex is stabilized via multiple protein:lipid interactions

The 3D reconstruction of the nanodisc-embedded complex reveals several rod-shaped densities proximate to PelBC (Figure 5). Within the periplasmic leaflet, most of those densities emerge from the N-termini of PelC subunits, so they are unambiguously assigned to the covalently bound acyl chain anchors. For several PelC subunits, all three acyl chains conjugated to the N-terminal cysteine are resolved, reaching up to 16 carbon atoms (Figure 6B and Suppl. Figure 4D). The anchor chains of individual PelC subunits are sandwiched between two Trp-149 residues, one from the same PelC, and one from a preceding subunit, a configuration that potentially facilitates docking of the lipoproteins at the membrane interface. Compared to the membrane-less crystal structure (Marmont *et al*., 2017a), Trp-149 residues undergo rotation towards the lipid bilayer (Figure 4A), so their indole rings immerse into the hydrophobic core of the membrane. Previous *in vivo* analysis showed that *P. aeruginosa* biofilm formation at the water-air interface was severely inhibited once Trp-149 was replaced by alanine (Marmont *et al*., 2017a). To test whether this functional defect arose from the complex assembly, we expressed PelC_W149A_ mutant either alone or in combination with PelB. PelC was targeted to the outer membrane in both cases, however we found a detrimental effect of the mutation on stability of the PelBC complex, as only PelB barrel without the lipoproteins could be isolated (Suppl. Figure 8). Thus, we concluded that Trp-149 stabilizes the lipid anchors within the assembled PelBC complex, possibly by suppressing their dynamics, and it may facilitate the oligomerization of the subunits into the ring.

**Figure 5.**
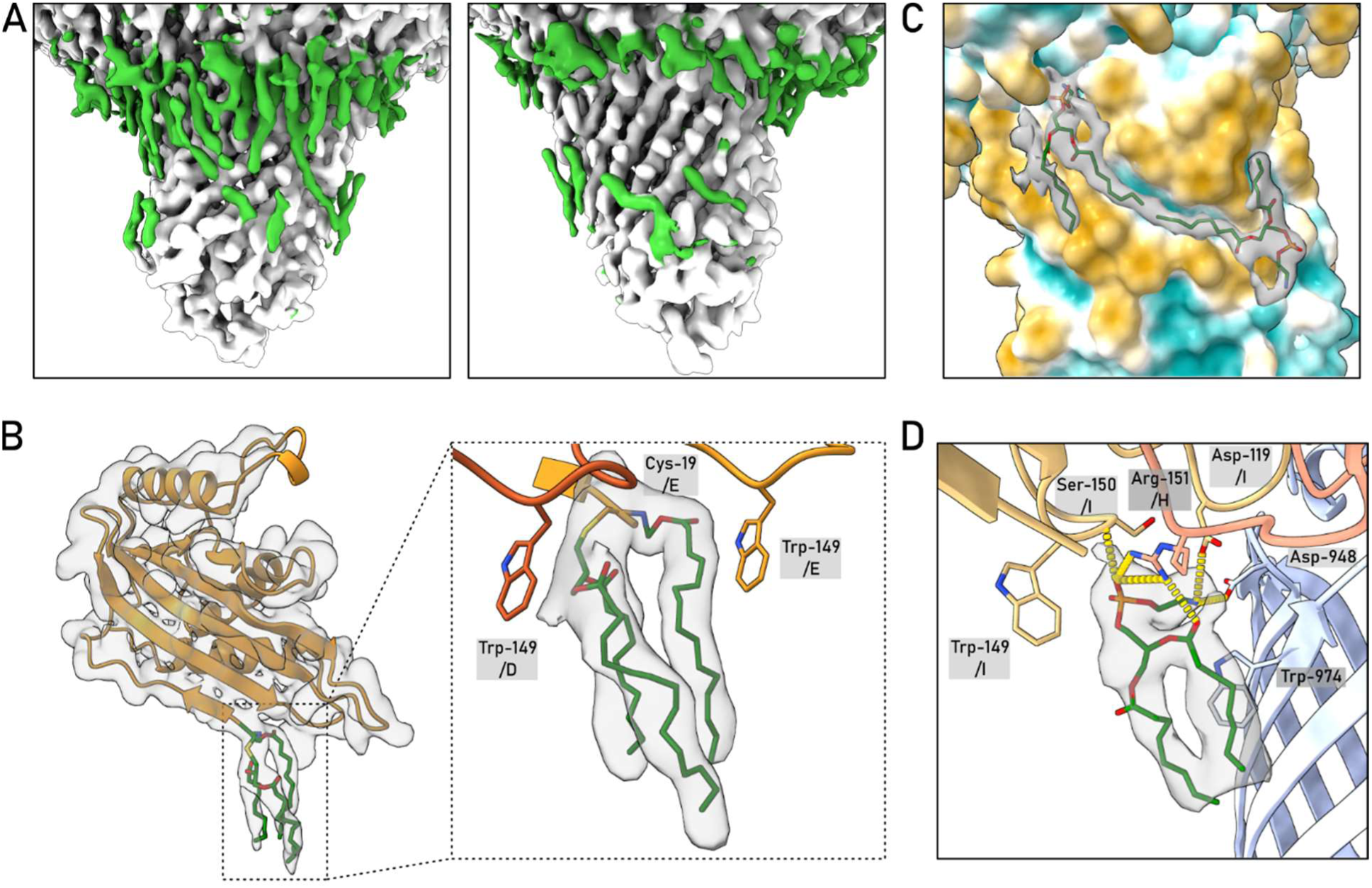
The proximate lipid environment of the PelBC complex. (A) Non-proteinaceous densities (green) within the nanodisc assigned to immobilized acyl chains of lipids and the lipoprotein anchors. (B) The N-terminal acyl chains of the lipoprotein PelC (on example of the subunit E) are stabilized by Trp-149 residues from two subunits, D and E. (C) The acyl chains are commonly docked within the hydrophobic grooves of the β-barrel. (D) Modelled structure and the interaction network of PE lipid molecule proximate to PelC subunits H and I.

**Figure 6.**
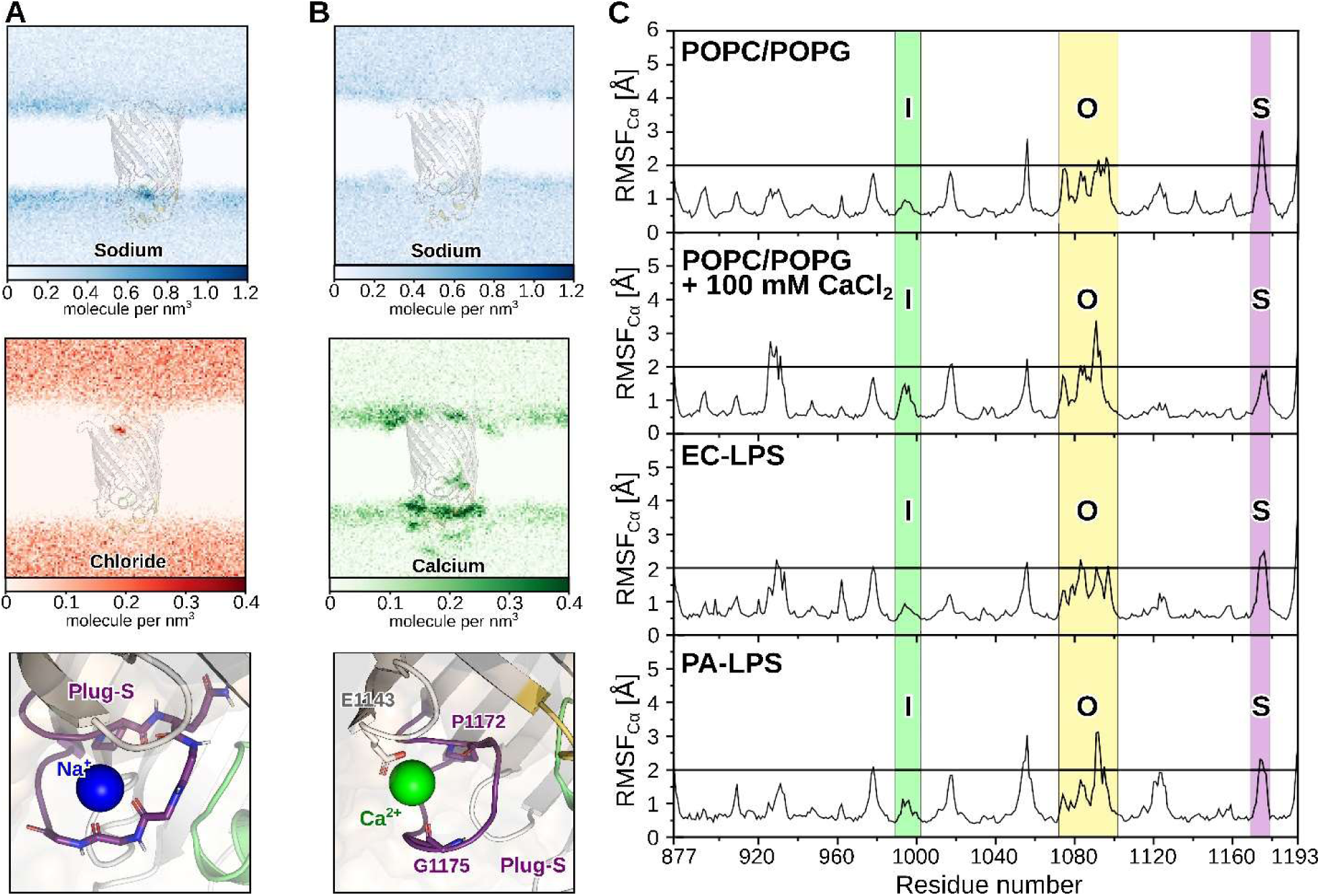
The ion distribution and the loop dynamics during the MD simulations. (A) The distribution density maps of sodium and chloride ions across the PelB β-barrel embedded into the symmetric POPC:POPG lipid bilayer. The Plug-S loop stably docks a sodium ion via interactions with the carbonyl groups (lower panel). (B) The density maps of sodium and calcium ions in the same settings as (A). A calcium ion binds stably at the side of the Plug-S via the Pro-1172, Gly-1175 and Glu-1143. (C) The dynamics of PelB residues depicted via the RMSF of the Cα atoms over the time of 500 ns in symmetric POPC/PPG lipid bilayer with and without calcium ions, as well as native-like membranes with an LPS leaflet based on *E. coli* and *P. aeruginosa* models in CHARMM-GUI. Residues with displacement over 2 Å are considered as flexible. The regions corresponding to Plug-I, Plug-O and Plug-S loops are highlighted. Representative simulations of each system are shown, while all plots of the triplicates are shown in **Supplemental Figures 11** and **12.**

The lipid anchors of PelC subunits reside at 3-4 Å distance from the PelB barrel, and so do not interact with the protein. However, multiple densities are found at the surface of PelB, which occupy the grooves formed by the membrane-facing aromatic and apolar residues (Figure 5A and C). Those were assigned to acyl chains of the phospholipids, which were either co-purified with the PelBC complex, or were provided upon the nanodisc reconstitution. A fully resolved lipid molecule was located between the PelC subunits H and I, forming extensive interactions with the lipoproteins and the barrel (Figure 5D). Based on the size and the interactions with the proximate polar residues which coordinate the head group, the density was assigned to a zwitterionic lipid, either PE or PC (Suppl. Figure 9). The best-fitting PE could be co-isolated from the bacterial membrane, while PC is a constituent of the synthetic nanodisc. Remarkably, a continuous density was resolved in a groove crossing the strands 1-2 and 13-16 that stapled N- and C-terminal ends of the β-barrel. The density spanning from the periplasmic to the extracellular sides of PelB likely arises from averaging of acyl chains that belong to lipids in the opposing leaflets. In the native bacterial membrane, the groove would be partially filled by a lipid A molecule in the extracellular leaflet. In support of this hypothesis, a cationic cluster of Lys-888, Lys-897 and Arg-921 is found at the extracellular side of the groove, offering a docking site for the phosphate head groups. On the other side of the barrel, a similar cluster of Arg-1071 (β-strand 11) and Arg-1102 (β-strand 12) is formed near a hydrophobic groove, which appears to be well-conserved feature among *Pseudomonas* species (Suppl. Figure 5), suggesting that the PelB structure is evolutionary tuned to facilitate interactions with lipid A molecules.

### In silico molecular dynamics of PelB in the lipid membrane

The cryo-EM analysis revealed a single conformation of PelB, where the groove within the periplasmic TPR’s is aligned with the pore at the extracellular side of PelB, thus forming a route for the polysaccharide. However, the exit pore of PelB between the strands 12-16 and the opposing Plug-I is occluded by the glycine-rich loop Plug-S (Figure 3). Analysis of the tunnels available within the resolved PelB structure shows that the width at the bottleneck is just 4 Å (Figure 3B). This tunnel cannot be employed for the polysaccharide translocation requiring a width of ∼8 Å (Morgan *et al*, 2013), so a conformational change, such as dislocation of Plug-S, must occur to open the conduit. Notably, Tyr-1102 in the proximity of Plug-S is universally conserved among PelB homologs, occasionally being substituted by phenylalanine and also found in the BcsC structure, so it may be involved in the EPS translocation, e.g. via CH-pi interactions with the pyranoside rings. We speculated that the PelB conformation, first of all its extracellular exit, may be affected by multiple physiological factors, such as presence of divalent cations, LPS, and/or variations in the temperature. To test those, we established all-atomic simulations of the PelB β-barrel (residues 877-1193) under a set of relevant conditions, where the symmetric phospholipid bilayer was compared to two native-like membranes containing LPS molecules (Suppl. Figure 10). Equilibration of the nanodisc-like phospholipid-based system in presence of different ions revealed remarkable features at the extracellular side of PelB: The highly charged region consistently attracted cations, and Plug-S formed stable interactions with a de-solvated sodium ion coordinated by the carbonyl groups of the tetra-glycine loop (Figure 6A). Once CaCl_2_ was included in the simulation, a calcium cation displaced Na^+^ from Plug-S and was stably docked between Pro-1172, Gly-1175 and Glu-1143 from a proximate extracellular loop (Figure 6B). Despite the significant difference in the ion distribution, the dynamics of the protein were similar between the tested conditions, where Plug-O and Plug-S repeatedly crossed the threshold of 2 Å, while Plug-I remained rather static (Figure 6C and Suppl. Figure 11).

The mimetics of *P. aeruginosa* and *E. coli* outer membranes available in CHARMM-GUI differ substantially in their phospholipid composition in the periplasmic leaflet, but also the length of the O-antigen polysaccharides of LPS, which extend into the solvent by 2.5 and 10 nm, respectively (Suppl. Figure 10). As PelB Plug-O is exposed to the LPS environment, its interactions were analyzed on the example of *E. coli*-type membrane by measuring the average distance between the exposed residues and the specific regions of the extended LPS (Figure 7A). The interactions were clustered around three arginines reaching the phosphate groups of the lipid A (Arg-1071 and Arg-1102) and the inner core (conserved Arg-1075). As a result, the latter conserved cationic site attracted two LPS molecules, and two other LPS were docked by Lys-888, Lys-897 and Arg-921, as predicted from the cryo-EM structure (Figure 7B and C). However, no consistent deviations in PelB RMSF values were observed between the simulations in either *E. coli* or *P. aeruginosa* outer membrane models (Figure 6C and Suppl. Figure 12). The experiments were performed at 25 °C, a relevant condition for the EPS secretion in surface-based biofilms. To test whether the protein dynamics, such as a movement of Plug-O observed in a single trial (Suppl. Figure 12), are enhanced by the temperature, we repeated the simulations at 37 °C. Remarkably, the elevated temperature favored further dislocation of Plug-S, especially for PelB in *P. aeruginosa* membrane (Suppl. Figure 12) and individual conformations of PelB visualize transient pores, which are wide enough for passage of multiple water molecules (Supp. Figure 13). The position of the loop within the PelB structure, its high dynamics and the prominent interactions with cations together suggest that Plug-S is the actual gating element for translocation of the cationic polysaccharide.

### Single-channel conductivity reveals conformational dynamics of PelB

As opening of PelB is required only once the cationic polysaccharide is encountered, a change in the electrostatic environment might be a potent trigger for switching the protein conformation. To probe the PelB dynamics under the altered electrostatic conditions, we established single-channel electrophysiology experiments to measure the ion conductivity across the β-barrel. In the electrophysiology set-up, the protein of interest is embedded into a free-standing lipid bilayer separating two compartments, and the ion passage across the membrane is recorded upon applying an electrostatic potential. The conformational dynamics of channel-forming membrane proteins, such as ion channels, toxins, but also outer membrane porins result in the ion flow fluctuations (Willems *et al*, 2017), which can be then used to describe the protein dynamics in the physiologically relevant environment.

To investigate PelB gating, the protein was expressed in *E. coli* C43(DE3) *ΔacrAB ΔompF* strain to exclude potential contamination with the major porin OmpF (Kanonenberg *et al*., 2019). PelB insertion into the lipid bilayer of 1,2-diphytanoyl-sn-glycero-3-phosphatidylcholine could only be achieved at high ionic strength and low pH 3.5, which would compensate the anionic nature of the extramembrane loops. Upon applying a potential of –150 mV across the membrane, the inserted protein manifested mainly a closed state, but also transient events of higher conductance were observed, which indicated reversible switching to open conformations (Figure 9A). Due to stochastic bilayer incorporation and switching between the conformations, ion flow through multiple PelB proteins could be eventually measured, resulting in multiplex opening and closing patterns (Suppl. Figure 14). We used only single-channel traces for analysis of conductance and open/closing probabilities (Figure 9B and C, Suppl. Figure 15), based on the largest common denominating current. The recorded ion currents of ∼15 pA, ∼40 pA and ∼115 pA showed good consistency between individual PelB proteins and corresponded to the conductance levels of 0.1 ± 0.02 nS, 0.28 ± 0.07 nS and 0.74 ± 0.07 nS at −150 mV, 1 M NaCl. The identified conductive states are further referred as “closed”, “tunneled” and “open” conformations of the β-barrel, respectively. The closed state accounted for up to 45 % of measured time, and the rest was shared between the tunneled and open conformations. Each of the states was characterized by relatively high noise levels, which could originate from thermal fluctuations within the protein structure, first of all the dynamic Plug-S, while deletion of the periplasmic TPR domain did not affect the noise level (not shown). The upper conductivity level in the open state matches well with the values measured for OmpG of *E. coli*, 14-stranded porin with the wide central opening up to 1.5 nm (Liang & Tamm, 2007). Achieving this width in PelB would require a major re-arrangement of the sealing loops at the extracellular side. The conductivity of the tunneled state corresponded then to a channel of 0.8-1.0 nm, which is sufficient for the polysaccharide translocation (Morgan *et al*., 2013), and could be achieved by Plug-S dislocation, as suggested by transient conformations seen in MD simulations (Suppl. Figure 13).

**Figure 8:**
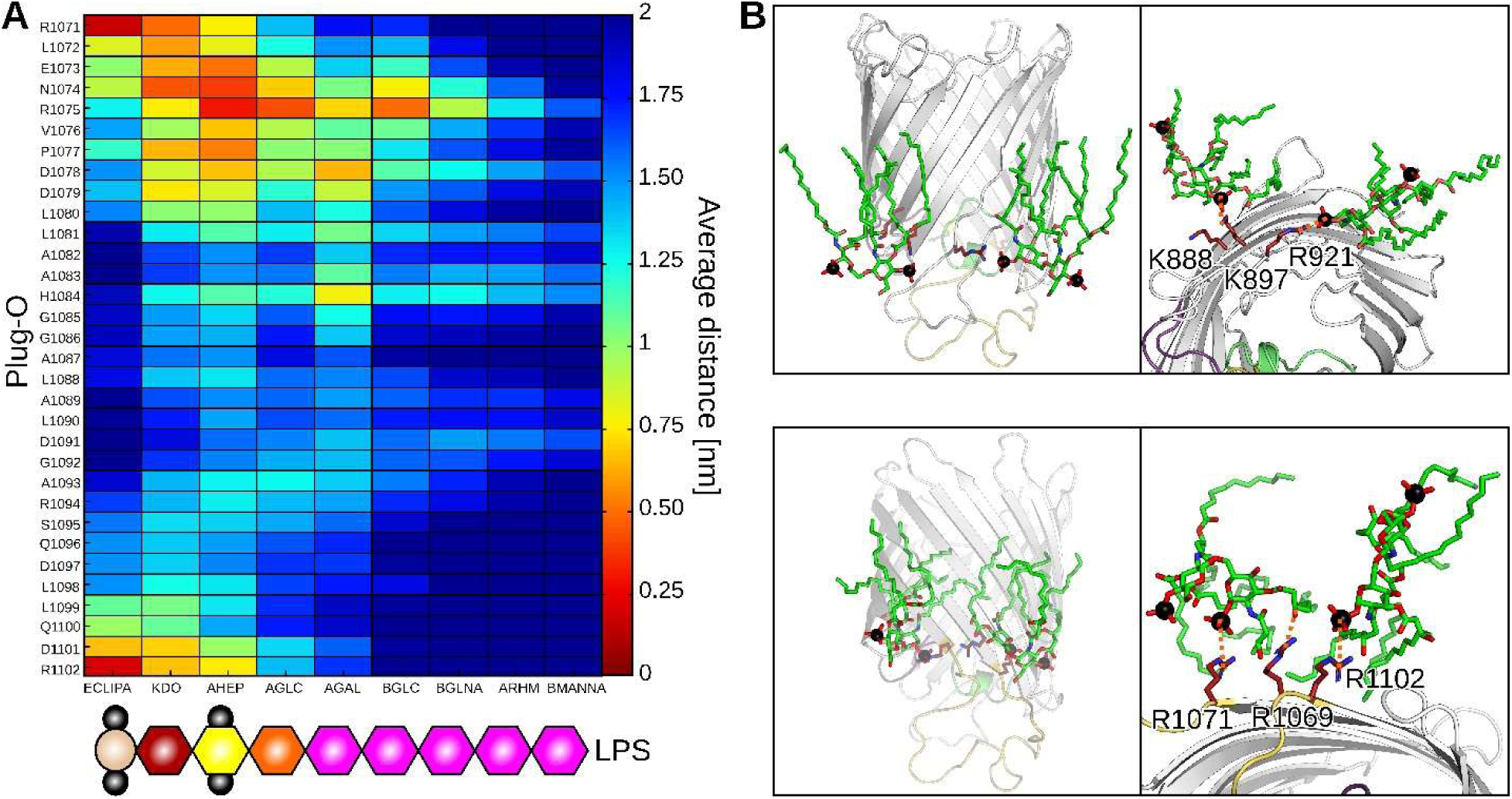
Interaction between LPS and the PelB β-barrel. **(A)** The average distance map between Plug-O and the LPS elements. The schematic structure of LPS molecule is shown below; details to the abbreviations are provided in the Suppl. Figure 10D. **(B)** A side and extracellular views of two interaction sides of PelB β-barrel with LPS. The key residues involved in the interactions with the phosphate groups of the lipid A or carbonyl groups of the fatty acids are indicated in the extracellular views.

**Figure 9.**
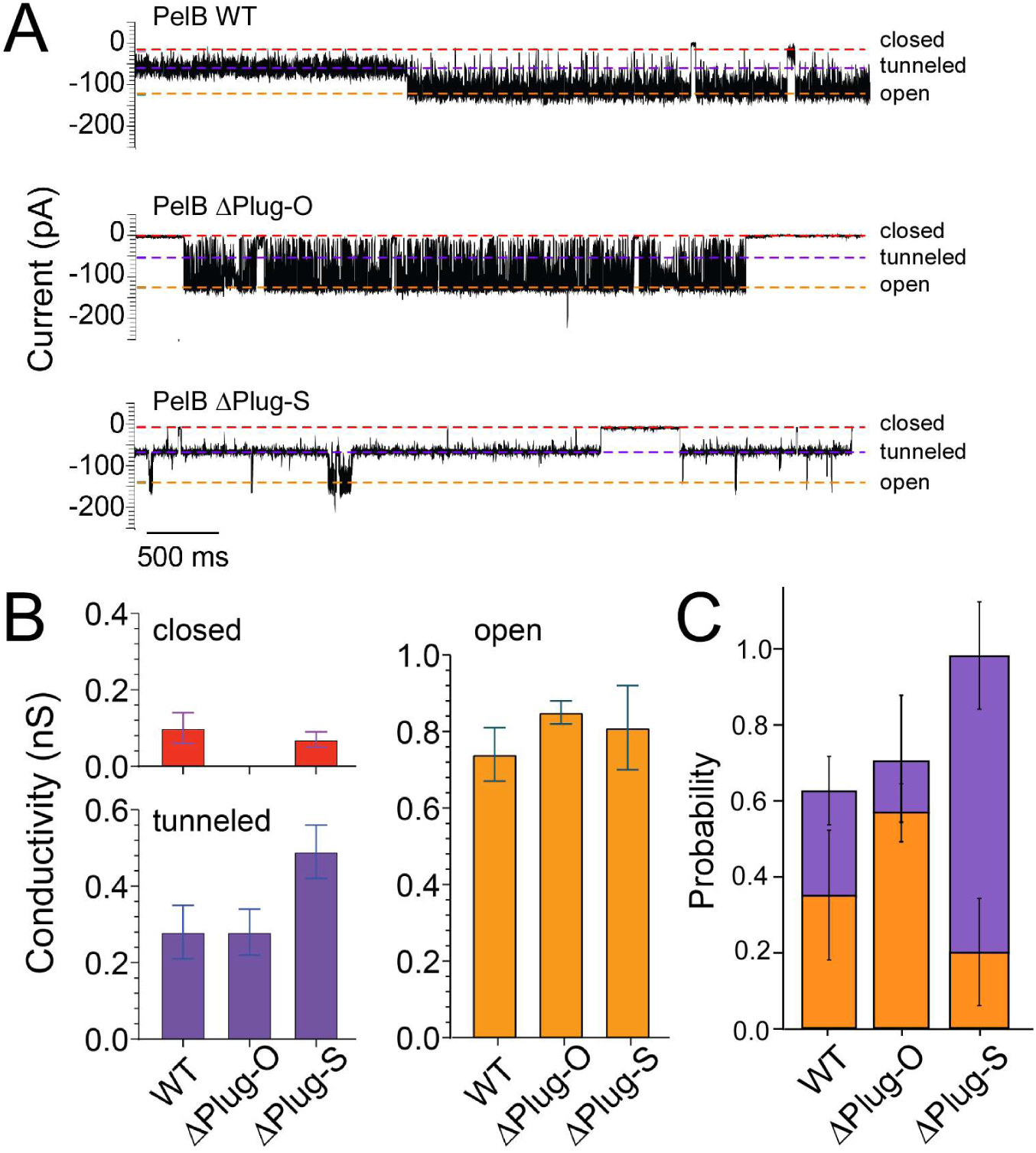
PelB dynamics in single-channel conductivity measurements. (A) Recordings of the ion current across individual PelB molecules, either wild-type (WT) or the truncated mutants ΔPlug-O and ΔPlug-S, suggest three conductance states to exist. (B) Calculated conductivity levels of each state measured for the PelB variants. (C) Probability of each conductance state determined for the PelB variants. Mean values +/- S.D. averaged from multiple recordings are plotted.

To probe the roles of the plug domains in the gating, PelB mutants with the individually deleted/shortened plugs were expressed and isolated for the conductivity experiments (Suppl. Figure 16). Among the mutants, only deletion of the Plug-I helical region (residues LMRAL) greatly reduced expression and purification yields of the protein, indicating defective folding. The detrimental effect of the Plug-I deletion was further proven when analyzing PelB stability via differential scanning fluorimetry (nanoDSF). When measuring the intrinsic fluorescence of the aromatic residues abundant within PelB (15 Trp within the β-barrel), we observed that the wild-type protein, as well as the Plug-O and Plug-S mutants underwent cooperative thermal denaturation in the range of 73-75°C (Suppl. Figure 16). In contrast, the construct lacking Plug-I unfolded already at 55°C, in agreement with the stabilizing role of the domain within the β-barrel structure, so the mutant was excluded from the further analysis.

Deletion of either Plug-O or Plug-S may affect the tunnel formation, as suggested by MD simulations and the tunnel calculations (Suppl. Figure 17). Indeed, in single-channel conductivity experiments, deletion of 14 residues within the non-conserved solvent-exposed loop Plug-O clearly favored the open state of PelB, however, both the closed and the tunneled states of the barrel were also observed (Figure 9 and Suppl. Figure 18). Once the Plug-S loop was removed, the closed state was nearly abolished (occurrence probability of 0.02 ± 0.02), and the protein existed predominantly in the tunneled conformation, occasionally switching to the high-conductance open state (Figure 9 and Suppl. Figure 19). Notably, the conductance of the tunneled state increased to 0.49 ± 0.07 nS and the noise level decreased, as the deletion of the flexible Plug-S widened the pore and eliminated the structural fluctuations (Suppl. Figure 17). The increased conductance value matched that measured for OmpC, where the pore width reaches 1.1 nm (Nikaido & Rosenberg, 1983). Thus, MD simulations and the conductivity measurements jointly suggest that the Plug-S dislocation within the wild-type PelB renders the tunneled state, which is sufficiently wide for the polysaccharide transport. The high-conductance open state may be a result of more extensive rearrangements, e.g. displacement/unfolding of the rigid Plug-I or deformation of the barrel under the applied potential.

## Discussion

Despite the abundance of biofilms in nature and the major roles they play in the microbial pathogenicity, the mechanisms of their assembly at the molecular level remain poorly understood. Resolving the structure and function of the cellular machineries involved in the biofilm formation, first of all exopolysaccharide synthesis and secretion, may assist in developing new strategies to suppress the spreading of many pathogens. Here, we focus on the mechanism of a unique secretion machinery for *P. aeruginosa* Pel exopolysaccharide and present the intact structure of the PelBC complex in the lipid environment, describe the determinants for its assembly, and provide the first experimental insights into the conformational dynamics that may facilitate translocation of the substrate exopolysaccharide.

Up to date, crystal structures of several exopolysaccharides transporters have been resolved, which mediate secretion of alginate (AlgE), PNAG (PgaA) and bacterial cellulose (BcsC) across the outer membrane ((Whitney *et al*, 2011); (Acheson *et al*., 2019); (Wang *et al*., 2016)). Consistently, they show β-barrels of either 18 or 16 strands occluded with loops at the extracellular side, and routes for the substrate translocation have been suggested based solely on the structural insights. PelB shows a high structural similarity to BcsC and PgaA, but as a unique feature in Pel secretion, the outer membrane barrel is essentially coupled to the ring of the lipoproteins PelC, which is fully resolved in our structure. The assembly is enabled by the conjugated acyl chains and the membrane-embedded Trp-149 of each PelC subunit, and the necessary electrostatic interactions of multiple lipoproteins with PelB, both with the helical TPR domains and the periplasm-exposed loops. It has been previously suggested that the negative charge at the central pore of PelC is required for electrostatic funneling of the cationic polysaccharide into the membrane channel. However, our data show that the key anionic residue Asp-119 within the D-loop is required for PelB:PelC docking, and the loop flexibility allows for building subunit-specific interactions at the interface within the asymmetric complex. Thus, the polysaccharide transport is rather dependent on the negative potential within the PelB barrel interior, i.e. the charged β-strands 11-14 and extracellular loops.

The structural analysis and the single-channel electrophysiology experiments suggest that PelB is a sealed β-barrel, not permeable in its idle state. Indeed, the outer membrane of P. aeruginosa is known to lack porins which would possibly allow passage of toxic agents across the membrane. Both the MD simulations of PelB in various environments and in vitro analysis of PelB mutants suggest that the channel opening required for the exopolysaccharide secretion is provided by a movement of the Plug-S, the extracellular loop aligned with the periplasmic TPR domains and the anionic wall inside the β-barrel. Remarkably, Plug-S is located in a direct proximity to Tyr-1103 from β-strand 12 (Figure 3C and Suppl. Figure 6), a residue which is highly conserved among PelB homologs from different species, including thermophiles, such as *Sulphurihydrogenibium azorense* and *Persephonella marina*, and also found in the cellulose transporter BcsC as Tyr-1030. The residue likely contributes to the channel and coordinates the polysaccharide via CH-pi interactions with pyranose rings. Here, further structural and biophysical analysis will be required to reconstruct the path of the Pel chain across the PelB barrel.

Based on the structure resolved in the lipid environment and the MD simulations in the native-like membranes we gained insights on the distribution of lipids and LPS molecules around PelB. While the apolar grooves formed at the outer barrel surface serve for docking the acyl chains, Arg/Lys clusters within the otherwise anionic extracellular interface facilitate local interactions with the phosphate groups of LPS. Recently, multiple studies highlighted the potential on amphipathic polymers for extraction of membrane proteins, followed by the structural and lipidomics analysis. On the other hand, computational modelling of the protein:lipid interactions, also in combination with cryo-EM data, allows identifying the interaction hot-spots and even the identity of the involved lipids (Ansell *et al*, 2023). Those approaches may be of the great value for studying further the PelBC complex, also in its native membranes of *Pseudomonas* species. Beyond the specific protein:lipid interaction, it remains to be shown whether the observed tilt of the PelC ring relatively to the PelB axis has an effect on the membrane morphology, e.g. increased curvature at the periplasmic side.

The necessity of the PelC ring and the importance of the involved PelC:lipid and PelC:PelB interactions remain puzzling. PelC subunits extensively interact with the TPR domains of PelB, which potentially serve as molecular rails for the Pel EPS through the periplasm, resembling the routes followed by polypeptide chains and lipid A in TamB and LptA transport proteins (Josts *et al*, 2017); (Suits *et al*, 2008). In the resolved structure, the groove of the TPR domain is aligned with the pore at the opposite side of PelB, with a broad electronegative surface in between, thus forming a continuous path for the cationic polysaccharide chain. The PelC assembly may assist in stable positioning of the TPR domains above PelB, though other transporters, BcsC and PgaA, equipped with similar TPR scaffolds do not require additional lipoproteins. Alternatively, PelC may facilitate the functionality of PelA, a bi-modal deacetylase/hydrolase enzyme of 105 kDa (Marmont *et al*., 2017b). PelA interacts with PelB residues 332-436 within the TRP region, where it can access the exopolysaccharide Pel and de-acetylate GalNAc moieties. The dimensions of the full-size PelA make it possible then to reach the PelC ring at the outer membrane, as it is predicted by the computational modelling (Suppl. Figure 20). As the high processivity of PelA may be required for efficient translocation of Pel, this additional interaction with PelC subunit K would stabilize the PelB-bound enzyme in the functional position. Testing this interaction *in vivo* and *in vitro*, also with help of structural analysis, should explain the necessity of this peculiar architecture and result in a comprehensive model of the exopolysaccharide export at the cellular boundary membrane.

## Acknowledgments

We would like to thank Susanne Rieder and Charlotte Ungewickell for the support with cryo-EM sample preparation and Prof. Dr. Albert Guskov (University of Groningen) for the initial cryo-EM analysis. We thank Prof. Dr. Birgit Strodel, Florian Altegoer and Sakshi Khosa (HHU Düsseldorf) and Jens Reiners (Center for Structural Studies, HHU Düsseldorf) for the fruitful discussions. We gratefully acknowledge the computing time granted through the Leibniz Supercomputing Centre (LRZ) of the Bavarian Academy of Sciences on the supercomputer SuperMUC-NG (project pn39gu). The hybrid computer cluster purchased from funding by DFG, project number INST 208/704-1 FUGG, and the Centre for Information and Media Technology at HHU Düsseldorf. The study was supported by the German Research Foundation (Deutsche Forschungsgemeinschaft, DFG; grant Ke1879/6 to A.K.), Dutch Research Council (Nederlandse Organisatie voor Wetenschappelijk Onderzoek, NWO; grant VI.C.192.068 to GM). RB acknowledges DFG Major Instrumentation grants for funding the cryo-EM infrastructure and the Advanced Grant from the European Research Council.

## Contributions

MB carried out sample preparations and biochemical analysis, and built the structural model based on the cryo-EM data. CRH curated the cryo-EM data collection and carried out the data analysis. SAPS carried out and analyzed single-channel conductivity experiments. JL performed molecular dynamics simulations. OB collected and curated the cryo-EM data. RB and GM curated the data analysis and secured funding. AK designed and coordinated the project, curated the data analysis and secured funding. All the authors wrote and edited the manuscript.

## Data availability

Coordinates and EM map have been deposited at the Protein Data Bank (PDB) under accession code XXX and the Electron Microscopy Data Bank (EMDB) under accession code EMD-XXXX.

## Methods

### Molecular cloning and protein expression

Gene sequence encoding *P. aeruginosa* PAO1 PelB (PA3063) residues 762-1193 with the conventional N-terminal pectate lyase signal peptide and an octa-histidine tag was synthesized by GenScript (Leiden, Netherlands). The *pelC* gene (PA3062) with the signal peptide of *E. coli* Lpp was synthesized by BioCat GmbH (Heidelberg, Germany). These synthesized genes were subsequently cloned into pETDuet-1 vector, *pelC* in MCS1 (restriction sites NcoI/HindIII) and *pelB* in MCS2 (NdeI/KpnI). For expression of PelB variants, pETDuet-1-based plasmids contained only the respective *pelB* genes. To introduce deletions in loops of the PelB β-barrel, the plasmid was amplified by PCR excluding the target DNA fragment and the template DNA was eliminated upon treatment with DpnI. Subsequently, the purified DNA underwent phosphorylation using T4 polynucleotide kinase and ligation employing T4 ligase. Molecular cloning was performed using chemically competent *E. coli* DH5α cells. All enzymes were from New England Biolabs. DNA isolation was performed with PCR & Gel Clean-Up and NucleoSpin Plasmid isolation kits (Macherey-Nagel). The sequences of the recombinant plasmids were confirmed through sequencing (Eurofins Genomics).

*E. coli* C41(DE3) *ΔompF ΔacrAB* cells (Kanonenberg *et al*., 2019) harbouring the recombinant plasmids were cultured in Luria Bertoni (LB) medium containing 100 µg/mL ampicillin at 37 °C with shaking at 180 rpm. Upon reaching optical density at 600 nm (OD_600_) of 0.6 to 0.8, protein expression was induced by addition of 0.5 mM isopropyl-β-thiogalactopyranoside (IPTG), followed by incubation for 3 hours at 30 °C. Cells were harvested by centrifugation at 5,000xg for 15 min at 4 °C. The cell pellets were resuspended in 20 mL resuspension buffer (40 mM HEPES pH 7.4, 150 mM NaCl, 5% glycerol). 0.2 mM AEBSF as well as DNase were added prior to the cell lysis step. Cells were lysed using a shear force (LM20 microfluidizer, Microfluidics), followed by centrifugation at 20,000xg for 15 min at 4 °C to remove the debris. To isolate the total membrane fraction, the clarified lysate underwent a further centrifugation step at 205,000xg for 1 hour at 4 °C (60Ti rotor, Beckman Coulter). The membrane pellet was then resuspended in the resuspension buffer, shock-frozen and stored at −80 °C until further use.

The localization of the PelBC complex in the outer membrane was verified through ultracentrifugation in sucrose gradient. The total membrane extracts were loaded onto a continuous 20-70% sucrose gradient (w/v) in the resuspension buffer prepared using an automated Gradient Station (BioComp Instruments). Subsequently, the gradient was centrifuged at 107,000xg for 16 hours at 4 °C (SW 40 rotor, Beckman Coulter). After centrifugation, the gradient was fractionated from top to bottom using the Gradient Station combined with the UV spectrometer and the fractions were analyzed by SDS-PAGE.

### PelBC isolation and reconstitution into nanodiscs

For isolation of the PelBC complexes and the variants of PelB, the total membranes were solubilized with 2% DDM (Glycon Biochemicals GmbH) in IMAC buffer (50 mM HEPES pH 7.4, 300 mM KCl, 5% glycerol) in 10x volume relative to the membranes. After 90 min incubation, the non-solubilized material was removed by centrifugation at 21,000xg for 15 min at 4 °C. The soluble fraction was supplemented with 2 mM histidine and loaded onto a gravity-flow column packed with 400 µL pre-equilibrated Ni-NTA agarose beads (Protino, Macherey-Nagel). Binding was carried out for 90 min at 4 °C with rotation, the flow-through fraction was collected, and the resin was washed four times with 2 mL of the IMAC buffer supplemented with 20 mM histidine and 0.05 % DDM. The target protein complex was eluted with the IMAC buffer supplemented 200 mM histidine and 0.05 % DDM. Three elution fractions of 800 µL each were pooled, concentrated to 500 µL and injected onto Superdex 200 Increase 10/300 GL size-exclusion chromatography (SEC) column in SEC buffer (50 mM HEPES pH 7.4, 150 mM NaCl, 0.03% DDM) connected to AKTA go system (Cytiva). Elution fractions were analysed via SDS-PAGE, and the protein concentration was determined spectrophotometrically based on calculated extinction coefficients of 95,910 M^-1^*cm^-1^ for PelB and 497,430 M^-1^*cm^-1^ for PelBC assuming 1:12 stoichiometry. Stability of the isolated PelB variants was analysed by differential scanning fluorimetry (nanoDSF) using Prometheus NT48 instrument (NanoTemper Technologies GmbH). The thermal denaturation was assessed by monitoring the intrinsic fluorescence at 330 and 350 nm between 20 and 95 °C through a temperature ramp of 1.5 °C/min.

Nanodisc-forming protein MSP1D1 was expressed and isolated as described (Ritchie *et al*., 2009). A lipid mixture composed of POPC:POPG lipids (molar ratio 70:30, Avanti Polar Lipids) was prepared in chloroform, the solvent was evaporated, and the lipids were suspended to the final concentration of 5 mM in 50 mM HEPES pH 7.4 and 150 mM NaCl. The formed liposomes were extruded stepwise through 1 µm and 200 nm track-etch membranes (Whatman) and solubilized using 0.5% DDM for 15 min at 40 °C. Subsequently, the purified PelBC complex was mixed with MSP1D1 and lipids at the molar ratio of 1:4:500 followed by 20 min incubation on ice. After the incubation, pre-washed Bio-Beads SM2 sorbent (Bio-Rad Laboratories) was added and incubated overnight at 4 °C with rotation. The reconstitution reaction was loaded onto Superdex 200 Increase 10/300 GL column connected to AKTA Pure (Cytiva), fractionated in 50 mM HEPES pH 7.4 with 150 mM NaCl and analysed via SDS-PAGE.

### Cryo-EM sample preparation and data collection

Before plunge-freezing, PelBC nanodiscs were supplemented with (1H, 1H, 2H, 2H-perfluorooctyl)-β-D-maltopyranoside (Anatrace) to a final concentration of 0.03 % to promote random orientation of the particles, and 3.5 µL of the sample were applied to glow-discharged Quantifoil Au 300 mesh R2/2 grids with an additional 3 nm layer of carbon. After incubation for 45 s, the grids were blotted for 3 s and plunge-frozen in liquid ethane using Vitrobot Mark IV (Thermo Fisher Scientific). Data collection was performed at 300 keV using Titan Krios microscope equipped with a Falcon 4i direct electron detector and a SelectrisX Energy Filter (all Thermo Fisher Scientific) at a physical pixel size of 0.727 Â. Dose-fractioned movies were collected in a defocus range from −0.5 to 3.0 µm with a total dose of 60 e-per Â^2^ and fractionated in 60 frames to obtain a total dose of 1 e-per A^2^ per frame.

### Cryo-EM data processing and model building

Gain correction, movie alignment and summation of movie frames was performed using MotionCor2 (Zheng *et al*, 2017). Further processing was carried out in cryoSPARC v4.4 (Punjani *et al*, 2017). A total of 31,712 micrographs were selected and CTF parameters were estimated using PatchCTF. Particle picking was carried out by running a Blob Picker job on a subset of 4040 micrographs. An initial set of 2,345,182 particles were extracted with a box size of 360 pixels, binned four times. After 2D classification, selected 2D classes containing 584,172 particles were used in one round of ab-initio reconstruction, followed by multiple rounds of heterogeneous refinement, which resulted in one good class with 386,144 particles. The particles of this class were re-extracted, binned two times, and further refined by non-uniform refinement. Selected templates were used to pick particles on the remaining micrographs of the data set. After multiple rounds of 2D classification a set of 4,773,524 particles was selected for downstream processing. The particles were sorted by heterogeneous refinement, using the refined volume and the three additional volumes from the ab-initio job. After multiple rounds of heterogeneous refinement, 3,174,321 particles converged to one good class, which was subsequently refined by non-uniform refinement. To further improve the resolution of the PelB β-barrel, masked 3D classification was performed and particles belonging to the classes with best resolved features were combined in a non-uniform refinement job (613,225 particles). Further 3D classification was carried out to improve the N-terminal TPR domain of PelB. A final set of 124,181 particles that converged to the best-resolved class was extracted without binning and refined by a non-uniform refinement job to a final resolution of 2.52 Å.

The complex model began with an AlphaFold2 prediction, which was fitted into the final cryo-EM volume. Subsequent manual adjustments were performed using the COOT program, with ongoing refinement achieved through the option real-space refinement in the PHENIX program. To model the PelC anchor chains, the fatty acid tails (and the glycerol backbone) of the phosphatidylethanolamine structure (PDB: PTY) were extracted and fitted into the density. The subunits of PelC were labelled from A to L, starting with the subunit closest to the first and last β-strand of the PelB barrel.

### Single-channel conductivity recordings

An electrophysiology chamber composed of two 500 µL compartments (*cis* and *trans*) separated by a 20 μm PTFE film with a central aperture of ∼100 µm was used for all experiments. To make a lipid bilayer, a drop of hexadecane (4% v/v in pentane) or a drop of hexadecane (4% v/v in pentane) was loaded on the *trans-*-side of the PTFE film, and allowed to evaporate for ∼2 min. Each compartment was then filled with 400 µL buffer (1 M NaCl, 20 mM citric acid pH 3.4) and two drops of lipids (DPhPC, 5 mg/mL). Ag/AgCl electrodes were inserted to each compartment: *trans* was the connecting electrode, *cis* was the ground electrode. By lowering and raising the buffer level in one compartment above the aperture, a lipid bilayer could be formed. The bilayer was equilibrated for 5-10 min before PelB was added. PelB samples were diluted 100-5000x with 15 mM Tris-HCl pH 7.5, 150 mM NaCl, 0.02% DDM and then added to the *cis* chamber in a small volume (<0.1 – 0.3 µL). Generally, the protein would insert upon applying –150 mV potential, and breaking/reforming the bilayer promoted insertion into the membrane. All experimental nanopore data were recorded under a negative applied potential (−150 mV), using an Axopatch 200B patch clamp amplifier connected to a DigiData 1440 A/D converter (Axon Instruments), and using Clampex 10.7 software (Molecular Devices, LLC). Data recordings were made in gap-free setting. Experiments were executed at least in triplicate. Measurements were conducted with a 50 kHz sampling frequency and a 10 kHz Bessel filter, unless otherwise specified.

Recordings were analyzed with Clampfit 10.7 software (Molecular Devices, LLC). Data was digitally filtered with a Gaussian low-pass filter with 1 kHz cut-off prior to analysis. Open pore current (I_open_), gating current (I_partially closed_), and closed current (I_closed_) were determined with Gaussian fit to all-point histogram with bin width of 0.5 pA. Only single channels were used for analysis. In most traces, porins entered the bilayer in multitude, and the single channel I_open_ was determined from the largest common denominating current. Trace parts with two or more porins were not used for analysis. The conductance (C, in nS) of open, partially open, and closed states was calculated from the corresponding Gaussian fit by

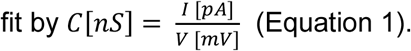

The probability (P) of open, partially open, and closed state was calculated by normalizing area under the curve (AUC) of the corresponding Gaussian fit to the sum of the AUCs, *e.g*.

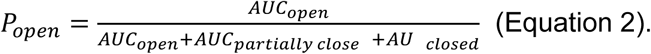

Per PelB variant, at least 3 technical replicates (N≥3 porins) of at least 8 s electrophysiology trace were used to calculate the average conductance and probabilities, in which the standard error of the mean (SEM) is reported in the error bars. Graphs were generated with GraphPad Prism software version 9.5.0.

### Molecular dynamics simulations

All MD simulations reported were performed on HILBERT from the Heinrich Heine University Düsseldorf or on SuperMUC-NG at Leibniz Supercomputing Centre in Munich. All systems are summarized in **Table 1** and in **Supplemental Figure 10A-C**. Two different LPS molecules with its components which are defined as the default for *E. coli* and *P. aeruginosa* were acquired from in CHARMM-GUI (Supplemental Figure 10D) (Jo *et al*, 2008); (Wu *et al*, 2014); (Lee *et al*, 2019). For more details on the following parts, a section is included in the **Supplementary information**. For all-atomistic MD simulations, the CHARMM36m force field (Huang & MacKerell, 2013); (Huang *et al*, 2017), modified TIP3P water and GROMACS 2018/2021 ((Abraham *et al*, 2015); (Lindahl, 2022b); (Lindahl, 2022a)) were accomplished while using a pressure of 1 bar. All systems were created by CHARMM-GUI and Ca^2+^ were added in the LPS layer. As protein structure, we used the cryo-EM structure of the PelB β-barrel (residues 877–1193) and the models of the truncated variants, PelB ΔPlug-O and PelB Δ ΔPlug-S (Mirdita *et al*￼￼(Abramson *et al*￼).

**Table 1.** Summary of simulations on PelB under different conditions.

| System and conditions | Size [atoms] | Runs | Cumulated time [μs] |
| --- | --- | --- | --- |
| EC-LPS, 150 mM KCl, 37°C | 184,012 | 3 × 500 ns | 1.5 |
| PA-LPS, 150 mM KCl, 37°C | 145,971 | 3 × 500 ns | 1.5 |
| EC-LPS, 150 mM NaCl, 25°C | 184,012 | 3 × 500 ns | 1.5 |
| PA-LPS, 150 mM NaCl, 25°C | 145,971 | 3 × 500 ns | 1.5 |
| POPC:POPG, 150 mM NaCl, 25°C | 105,713 | 3 × 500 ns | 1.5 |
| POPC:POPG, 150 mM NaCl + 100 mM CaCl<sub>2</sub>, 25°C | 105,707 | 3 × 500 ns | 1.5 |
| Plug-O mutant, POPC/POPG, 150 mM NaCl, 25°C | 102,598 | 3 × 500 ns | 1.5 |
| Plug-S mutant, POPC/POPG, 150 mM NaCl, 25°C | 104,249 | 3 × 500 ns | 1.5 |
| **Total** |  |  | 12.0 |

PA: *P. aeruginosa*; EC: *E. coli*; LPS: Lipopolysaccharide; PA-LPS: LPS with one O-antigen unit (outer leaflet) and 46% : 31% : 23% DOPE:DPPG:DPPE (inner leaflet) (Li *et al*, 2018). EC-LPS: 100% LPS with five O-antigen units (outer leaflet) and 75% : 20% : 5% POPE:POPG:PVCL (inner leaflet); 70% : 30% POPC:POPG (symmetric, used in experiments).

To check whether lipids and ions clustered within and around the PelB β-barrel in the in the *xz*-plane, we created density maps of ions and water via *gmx densmap*. Here, the direction to average was over *y*-axis, the grid size was set to 0.08 and the unit was molecule per nm^3^. We used the DuIvyTools (https://github.com/CharlesHahn/DuIvyTools) to change the format from *xpm* to a matrix *dat* file (*dit xpm2dat*), which could use for plotting the density maps via an own python script.

For describing the stability and flexibility of PelB barrel during the MD simulations, the root-mean-square fluctuations (RMSF) of the Cα around their average positions was calculated for each residue. The RMSF value represents the positional change of the selected atoms as time– average, where the residue with a value over 2 Å defines as flexible. To analyze PelB interactions, we generate average distance maps per residue between **(1)** the extracellular loops, as well as **(2)** to the ions and water, and **(3)** to LPS. We calculated the minimal distance (*gmx mindist*) between the groups by following computing the average distance over time.

To determine the geometry and the dimensions of the tunnel within the PelB β-barrel and its variants, HOLE algorithm (DOI: 10.1016/S0263-7855(97)00009-X) was employed in combination with the algorithm by MDanalysis (https://docs.mdanalysis.org/1.1.1/documentation_pages/analysis/hole2.html. The tunnels were visualized via VMD. To characterize the overall stability of the PelB barrel in the membrane, we further conducted clustering analyses using the algorithm of (Daura *et al*, 1999) as implemented in GROMACS. Here, the root-mean-square deviation (RMSD) is a measurement of the Cartesian deviation to a reference structure (mostly the start structure of the MD or a crystal structure). The clustering was employed for all atoms with a Cα-RMSD cutoff value of 0.15 nm by using the fitting function before calculating the RMSD on the structure to identify cluster membership. To generate the 3D protein structures, we used PyMol version 2.5. If not stated otherwise, analysis was computed with GROMACS. For plotting, Python or gnuplot were used.

## Supplementary materials

### Simulation details for the MD simulations of PelB barrel systems

All atomistic systems were created by CHARMM-GUI and the Ca^2+^ were added in the LPS layer. Furthermore, we used the *mdp* scripts for energy minimization, equilibration in five steps and production run from CHARMM-GUI web server since they are optimized for protein-membrane simulations. The energy was minimized to 1,000 kJ mol^-1^ nm^-1^ using the steepest descent algorithm, followed by five-step equilibration to the desired temperature of 298 K (25°C) or 310 K (37°C) and pressure of 1 atm to mimic the physiological environment. First, two *NVT* equilibration steps were applied to keep constant the number of atoms (*N*), the box volume (*V*), and temperature (*T*), followed by three-step *NpT* equilibration to adjust the pressure (*p*). The protein’s and lipid’s heavy atoms were restrained to allow the water molecules and ions to relax around the solute but they were decreased by every equilibration step. The Berendsen thermostat was employed to regulate the temperature in the *NVT* simulations, while the Berendsen thermostat and the semi–isotropic Berendsen barostat were employed for the *NpT* simulations. The PME method was applied to calculate long-range electrostatic interactions with periodic boundary conditions. The vdW and Coulombic interaction cutoffs were set to 1.2 nm using the LINCS algorithm to constrain all bond-lengths to hydrogens. Production MD runs were performed for 0.5 µs with a time step of 2 fs by recording the coordinates and velocities every 20 ps as well as the Nosé-Hoover thermostat and the semi–isotropic Parrinello-Rahman barostat.

### Supplementary tables

**Tabel S-M1:**
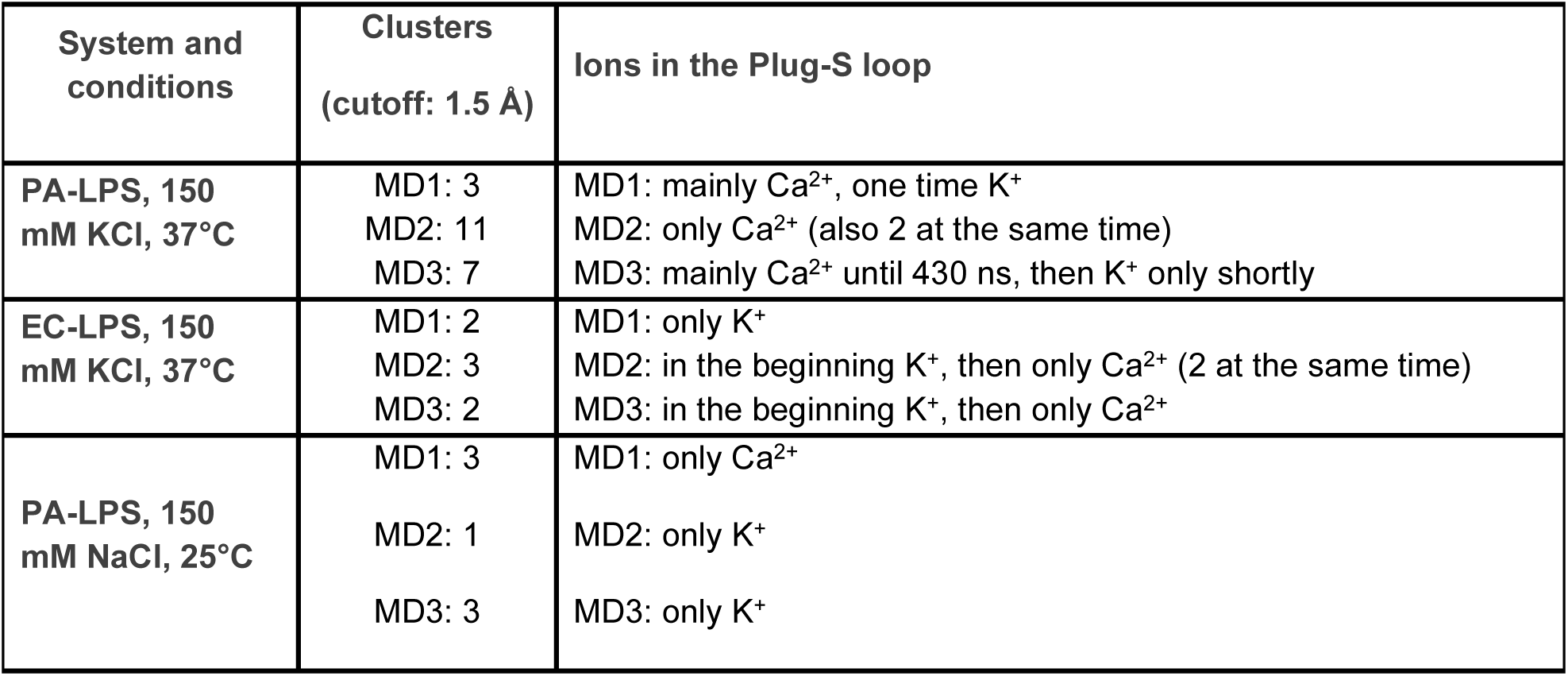
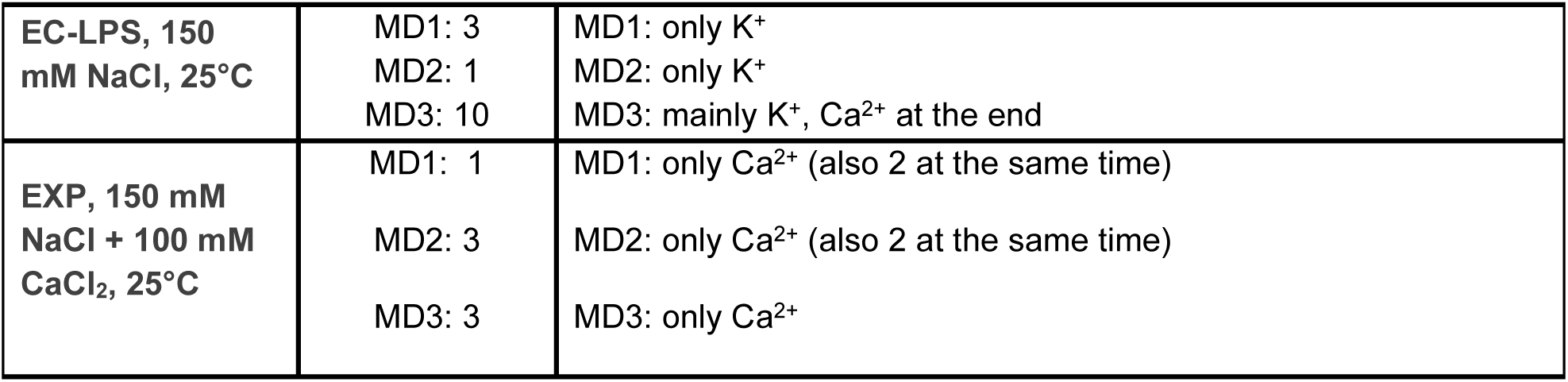
All simulations with Ca^2+^ ions used in the work with various statistical values. The cluster size means how many different structures with the cutoff of 1.5 Å were found. Furthermore, a description of the ion location at the Plug-S loop is given.

### Supplementary figures

**Supplemental Figure 1.**
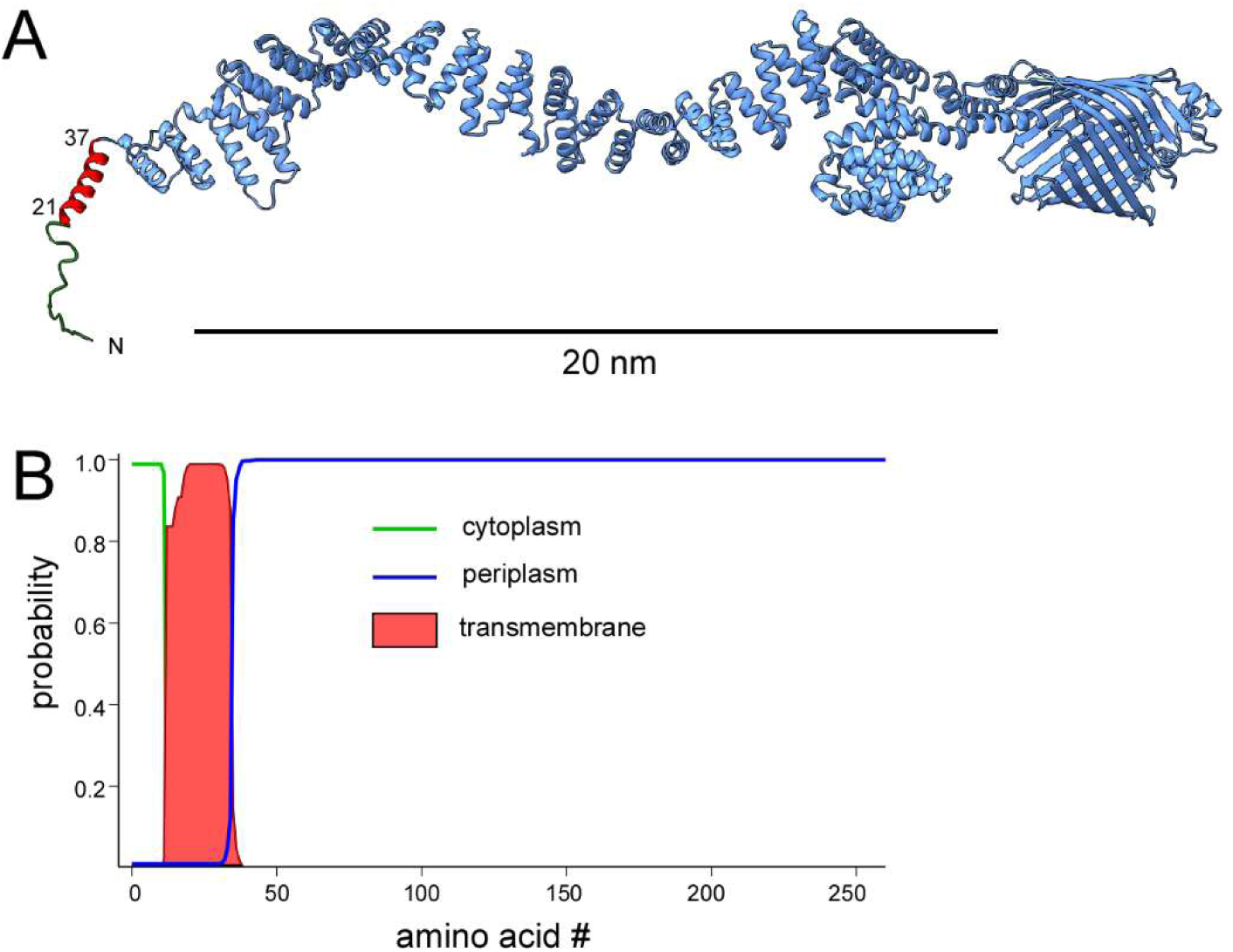
Outer membrane protein PelB is anchored in the inner membrane. (A) AlphaFold 3 model of *P. aeruginosa* PelB predicts an a-helix formed at the N-terminal end of the protein (shown in red) prior the periplasmic TPR domains. (B) TMHMM algorithm (https://services.healthtech.dtu.dk/services/TMHMM-2.0/) predicts a hydrophobic transmembrane helical domain within the same region of PelB, residues 21-38 (plot for the first 260 aa is shown). No signal peptide is identified using SignalP 6.0 service (https://services.healthtech.dtu.dk/services/SignalP-6.0/).

**Supplemental Figure 2.**
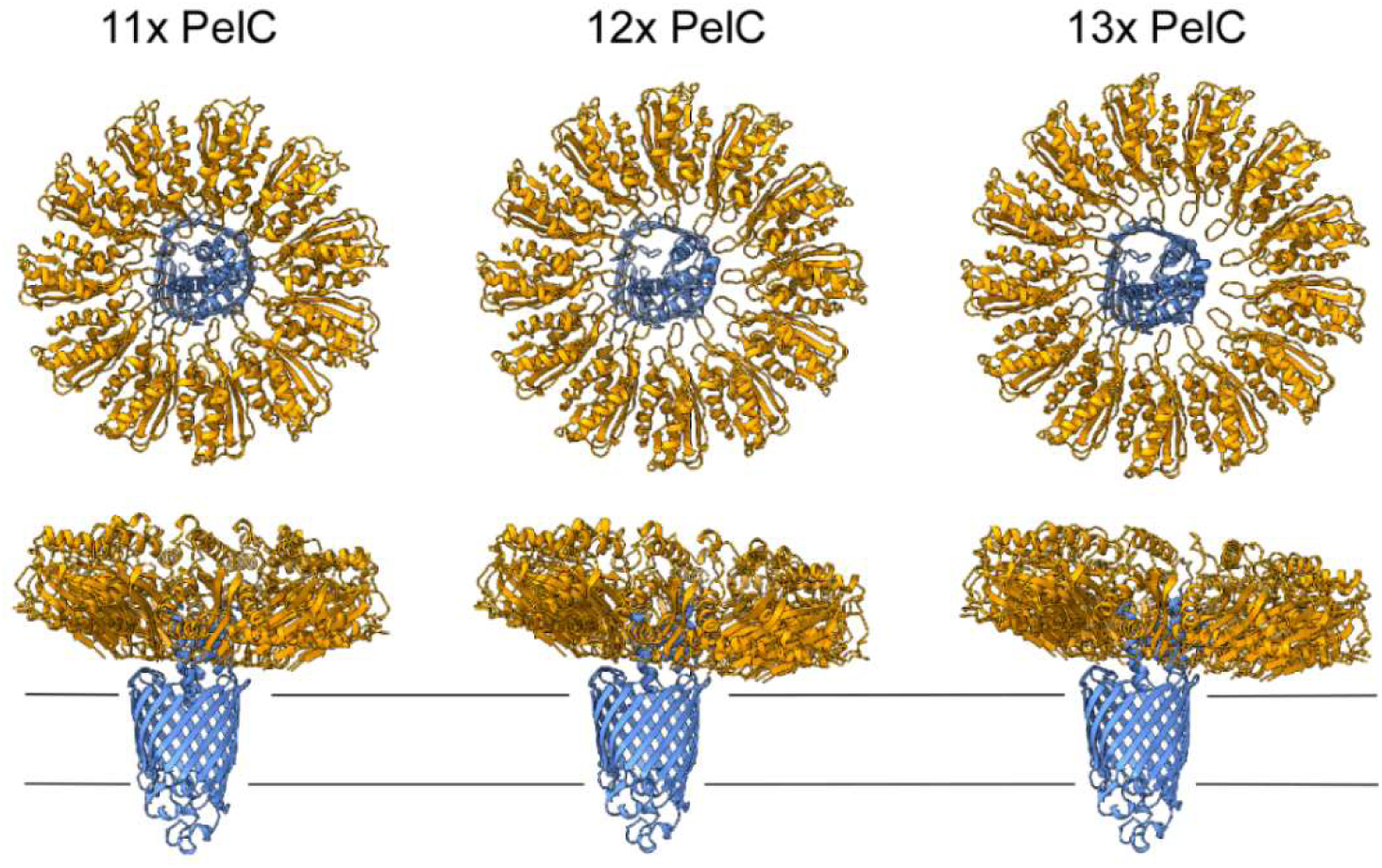
AlphaFold3 models of the PelBC complex assuming 11, 12 and 13 PelC subunits.

**Supplemental Figure 3.**
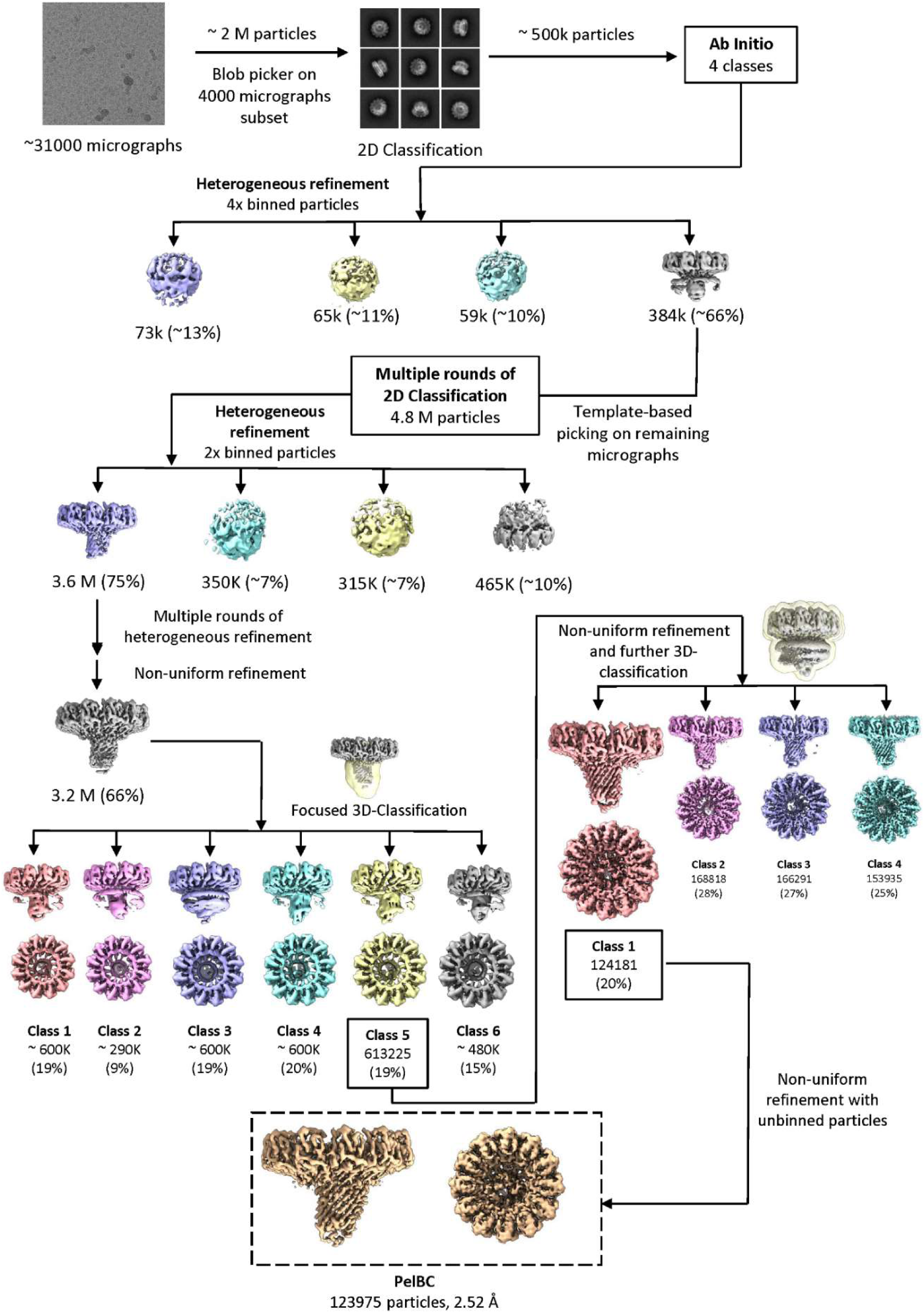
Single-particle analysis towards the PelBC structure. Summary of the sorting/refinement procedures performed upon single-particle analysis of the cryo-EM data set collected on the nanodisc-reconstituted PelBC complex.

**Supplemental Figure 4.**
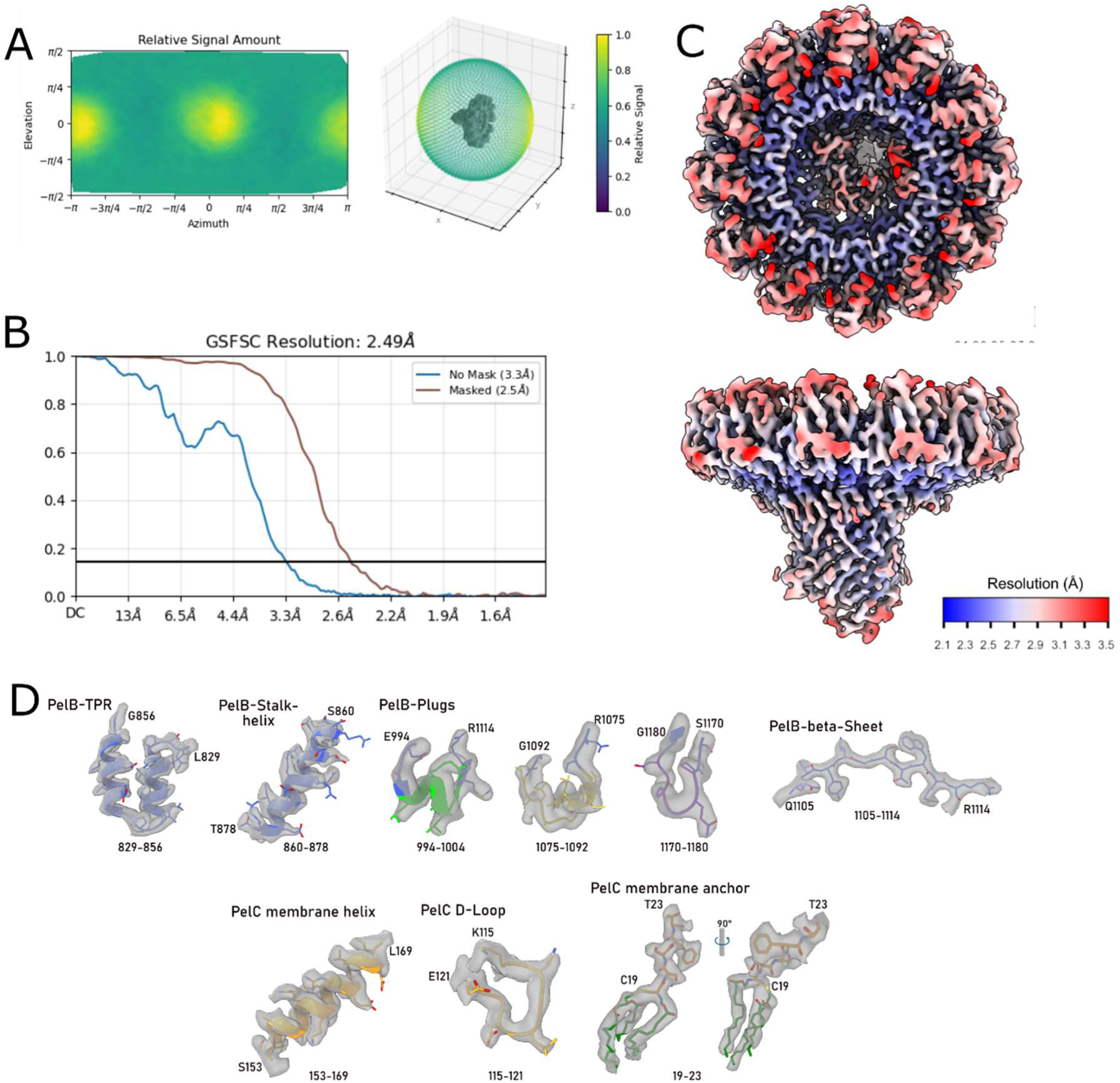
Global and local resolution of the PelBC map. (A) Angular distribution plot for the final reconstruction obtained from CryoSPARC. (B) Gold-standard Fourier shell correlation resolution curve for the final map, displaying resolution with a mask automatically generated by CryoSPARC. (C) Final PelBC map colored according to the local resolution as determined by CryoSPARC. (D) Selection of structural elements within the PelBC complex, with the corresponding densities from the cryo-EM reconstruction.

**Supplemental Figure 5.**
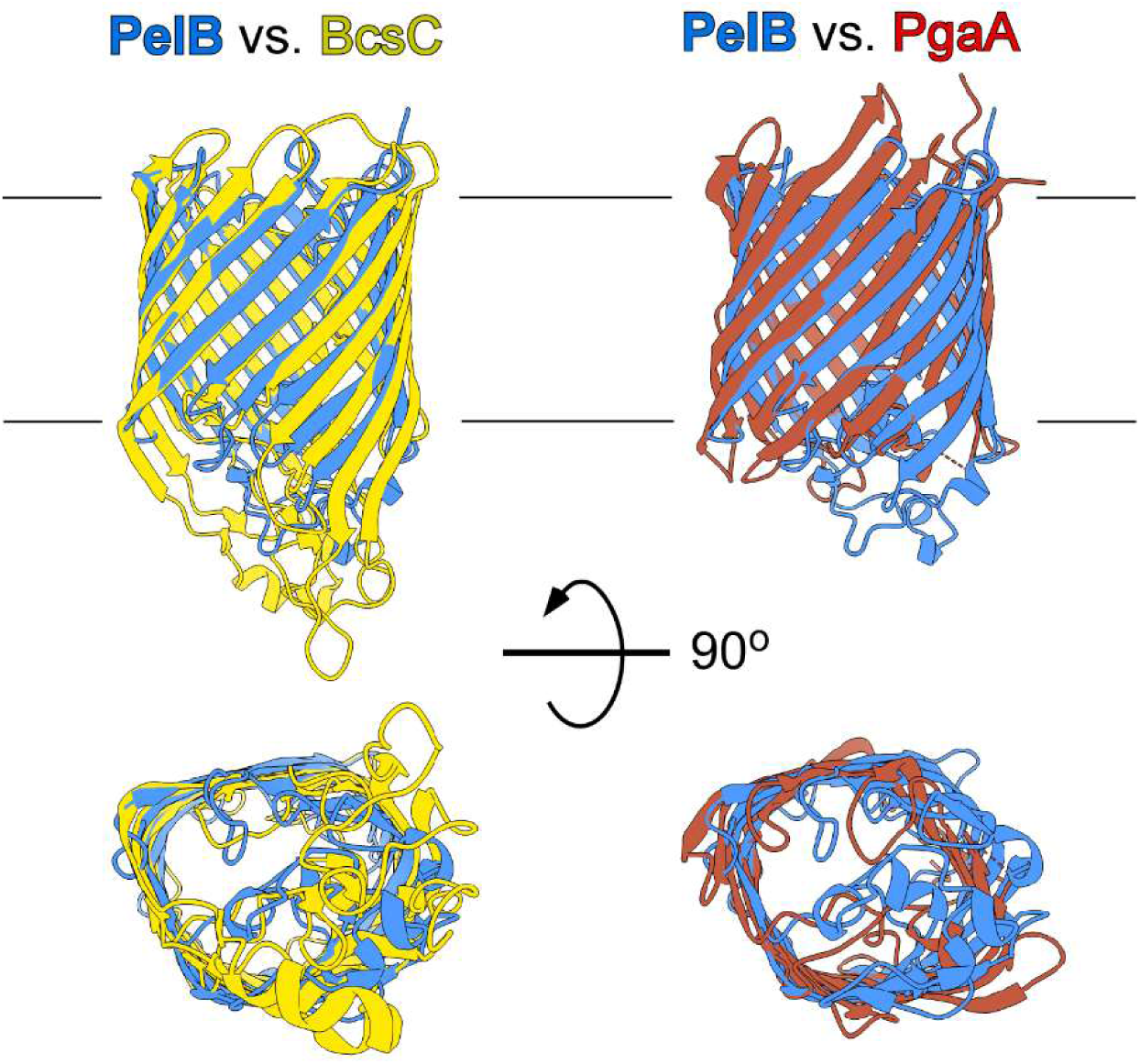
Conserved architecture of the bacterial outer membrane transporters of exopolysaccharides. Structure of *P. aeruginosa* PelB β-barrel superimposed with *E. coli* BcsC (PDB ID 6TZK) and *E. coli* PgaA (PDB ID 4Y25), views in the membrane plane and from the extracellular side.

**Supplemental Figure 6.**
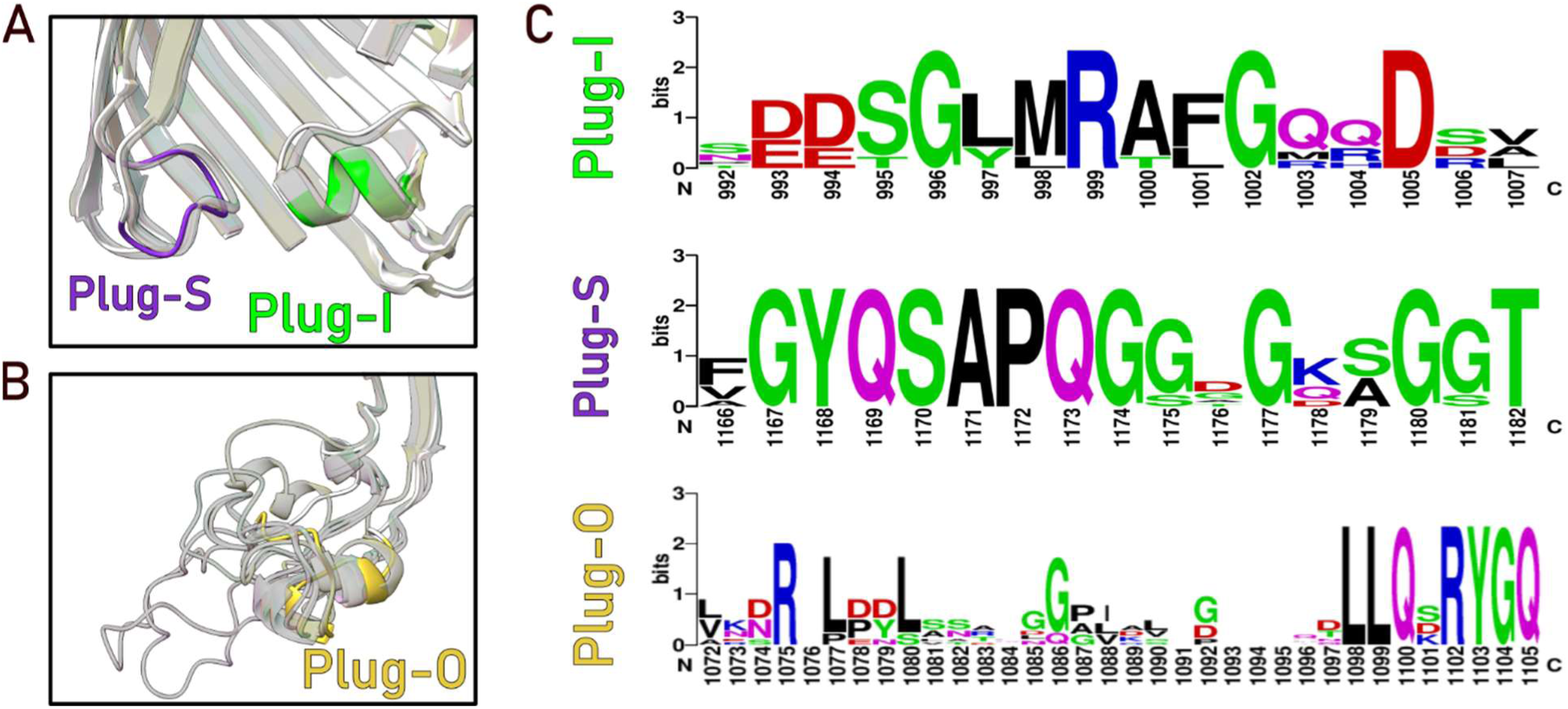
Conservation of the plug domains among *Pseudomonas* species (*P. aeruginosa, P. mandelii, P. oryzihabitans, P. protegens, P. simiae, P. sp., P. trivialis*).

**Supplemental Figure 7.**
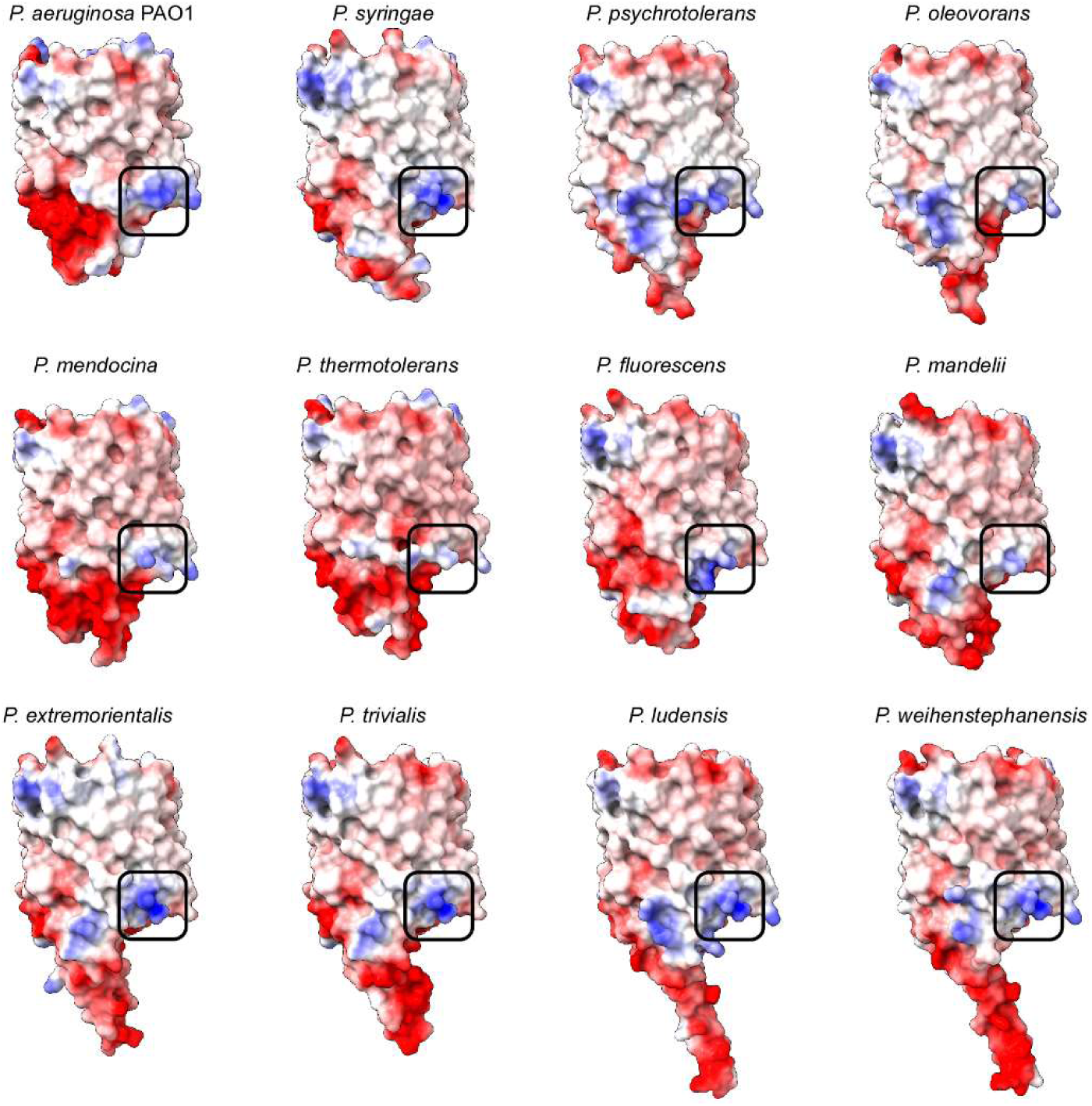
AlphaFold models of PelB homologs from the indicated *Pseudomonas* species highlight the conservation of the cationic site Arg-1071/Arg-1102 at the extracellular side of the β-barrel (black square) and the diversity of the exposed Plug-O domains.

**Supplemental Figure 8.**
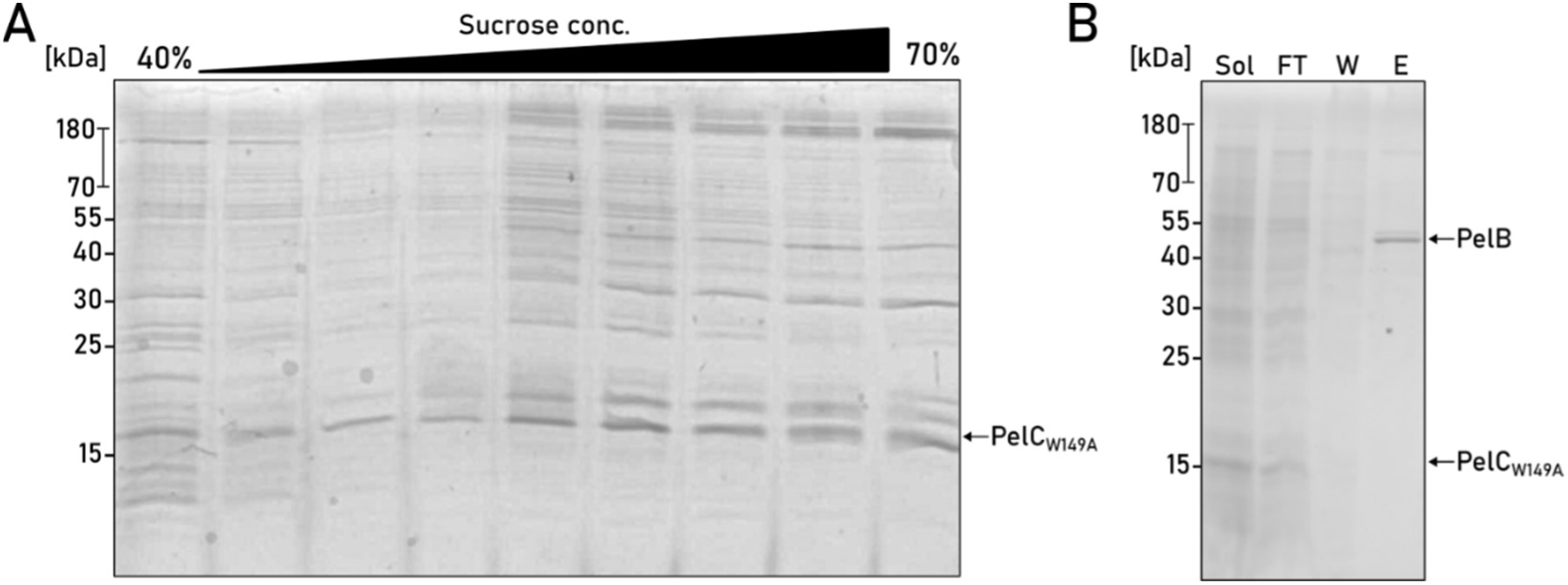
PelCW149A is targeted to the outer membrane but does not form stable complex with PelB. (A) SDS-PAGE of the sucrose density gradient shows localization of the PelC_W149A_ mutant in the late high density fractions corresponding to the outer membrane vesicles. (B) SDS-PAGE of IMAC purification of co-expressed PelB and PelC_W149A_ shows that the mutated PelC_W149A_ subunits are not co-purified with the His-tagged PelB (band). Loaded fractions: “Sol” - detergent-solubilized material; “FT” - flow-through; “W” - wash, “E” - elution.

**Supplemental Figure 9.**
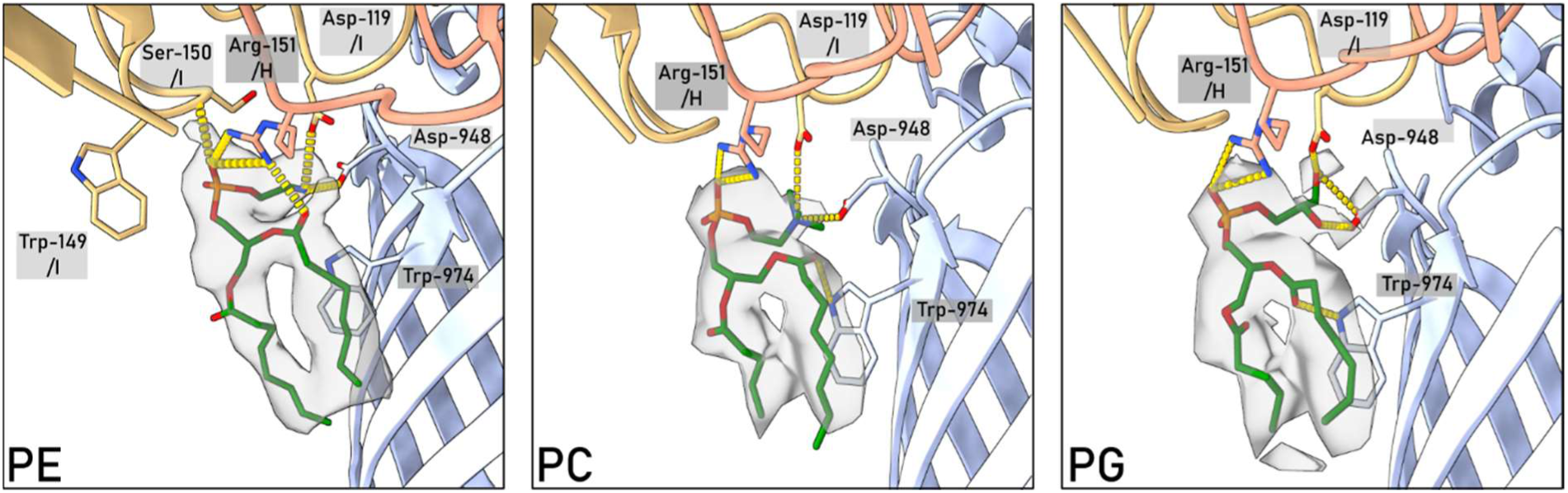
Phosphatidylethanolamine (PE) may form the most extensive interaction network with PelBC complex and ensure the optimal fit into the observed lipid density as compared to phosphatidylcholine (PC) and phosphatidylglycerol (PG).

**Supplemental Figure 10:**
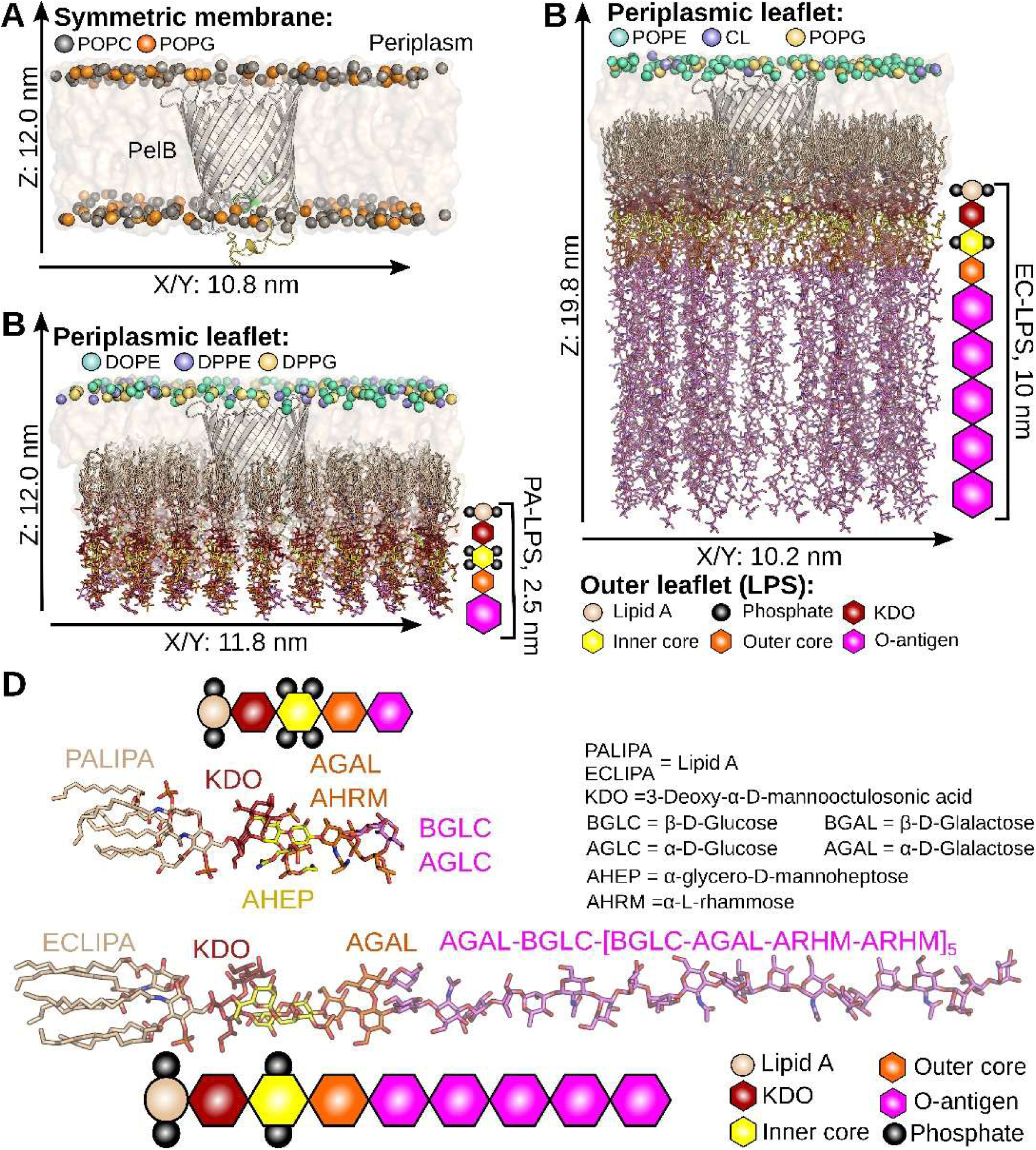
The PelB β-barrel systems for the MD simulations. **(A)** POPC:POPG system is shown like in the nanodisc experiments with 150 mM NaCl or additionally with 100 mM Ca^2+^ ions with 25°C as temperature. **(B, C)** CHARMM-GUI web server was used to model native-like membranes with LPS for *P. aeruginosa* (PA; inner leaflet DOPE:DOPG:DPPE). and *E. coli* (EC; inner leaflet POPE/POPG/CL) at 150 mM KCl and 150 mM NaCl as ion concentration, respectively. **(D)** The structure of the used LPS for *P. aeruginosa* (PA) and *E. coli* (EC) are depicted. In general, the LPS consists of the lipid A (wheat), which is the anchor in the membrane, the KDO (dark red), then sugar inner (yellow) and outer core (orange) as well as the repeated O-antigen units, where its length can be different for each organism. Phosphate groups are depicted as black circles at the sugars.

**Supplemental Figure 11.**
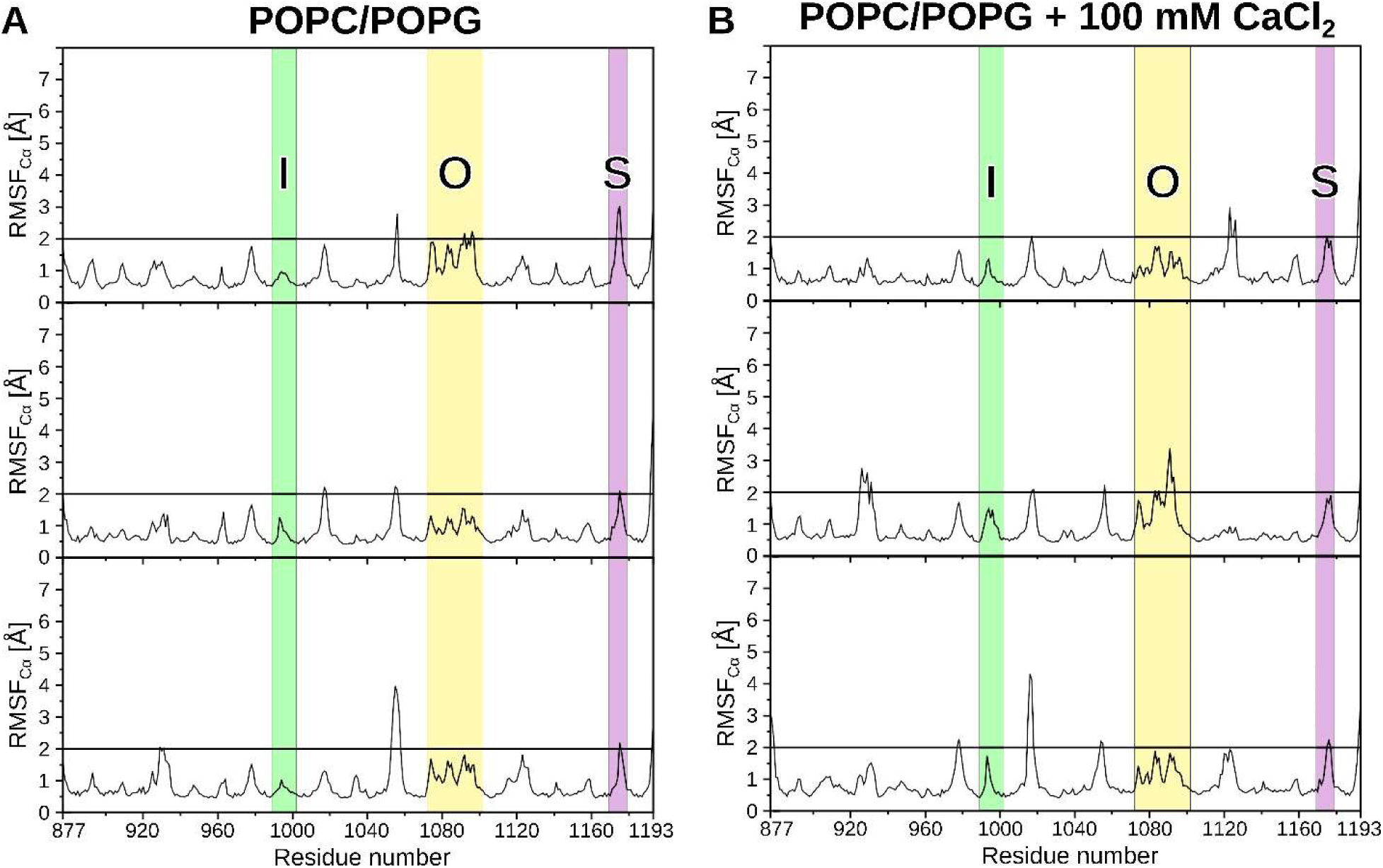
Dynamics of the PelB β-barrel in the POPC/POPG membrane system simulated in triplicates at 25°C and 150 mM NaCl without (**A**) and with 100 mM CaCl_2_ (**B**). To determine the flexibility, we used the root-mean square fluctuation (RMSF) of the Cα atoms per residue over the time, where residues over 2 Å are considered as flexible over 500 ns. The regions corresponding to Plug-I, Plug-O and Plug-S loops are highlighted.

**Supplemental Figure 12:**
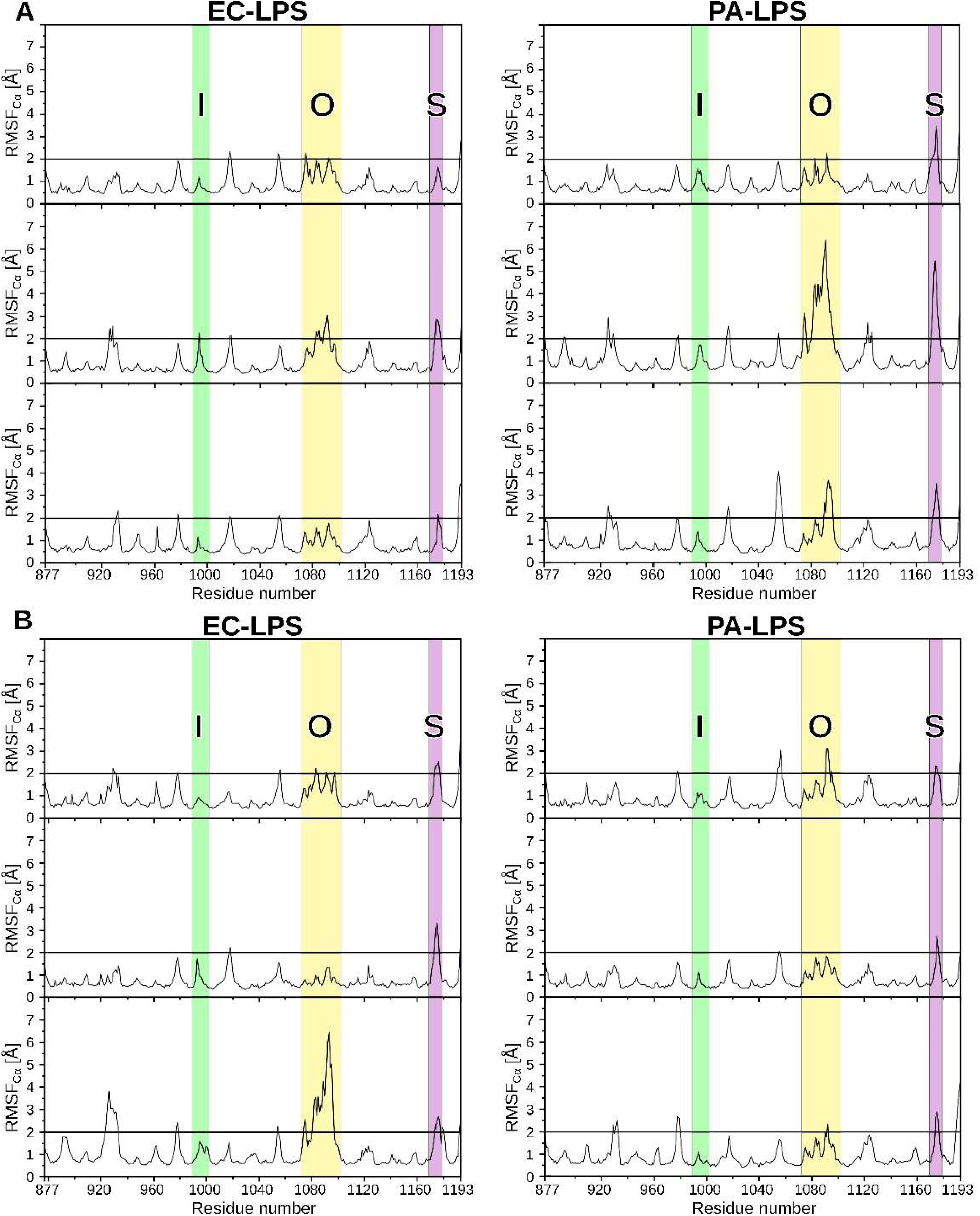
Dynamics of the Plug loops in the native-like LPS-containing membranes. To determine the flexibility, we used the root-mean square fluctuation (RMSF) of the Cα atoms per residue over the time, where residues over 2 Å are considered as flexible over 500 ns. Here, we used two conditions: **(A)** 37°C and 150 mM KCl and **(B)** 25°C and 150 mM NaCl. The regions corresponding to Plug-I, Plug-O and Plug-S loops are highlighted.

**Supplemental Figure 13.**
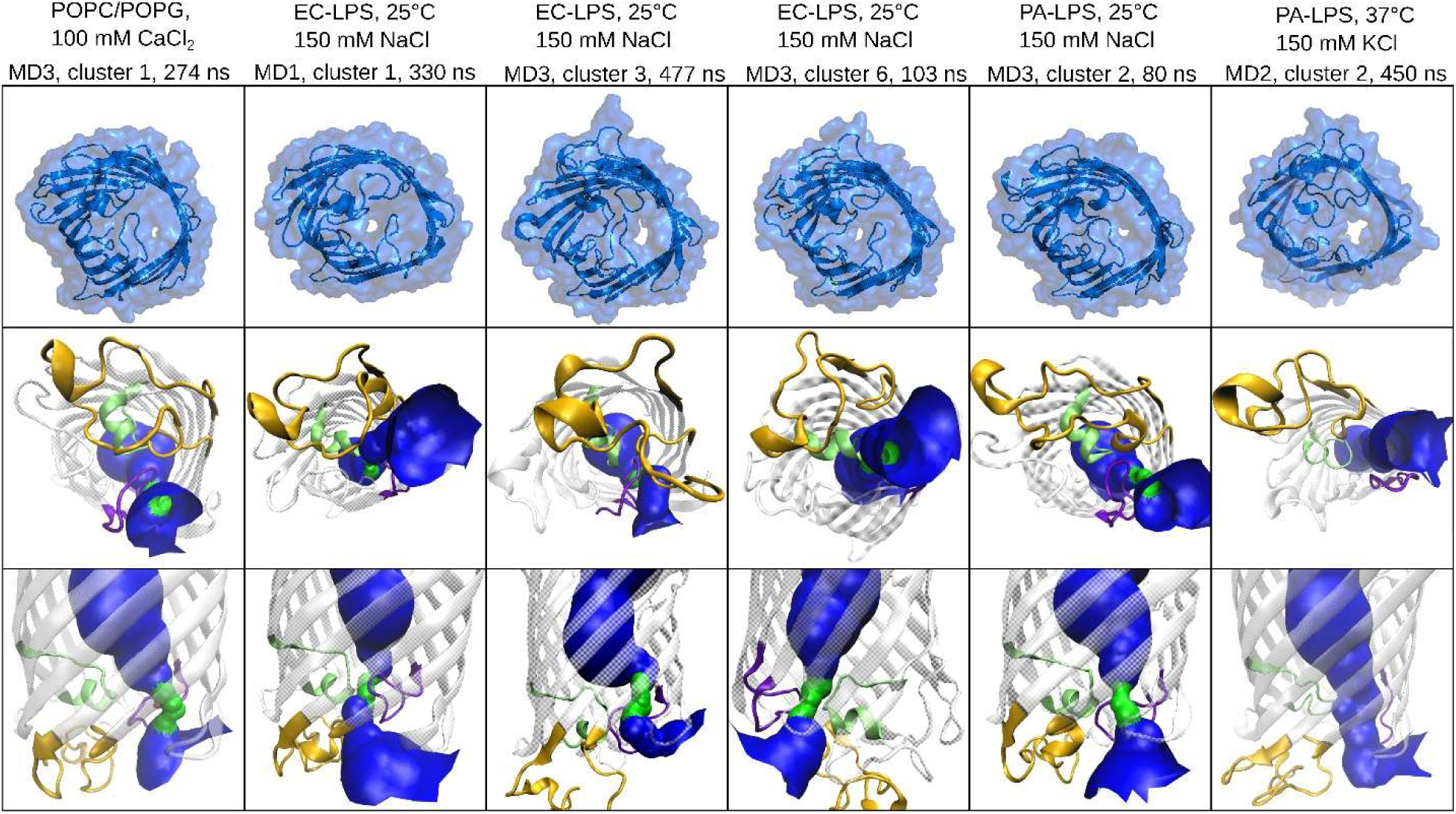
Transient pore opening in the wild-type PelB β-barrel in different membrane systems. The cartoon representation represents the orientation of the loops. The tunnel is colored in red for no water, in green for one water and blue for more water molecules, which are fitting in the pore. The pore analysis was done with HOLE algorithm and python package MDanalyis.

**Supplemental Figure 14.**
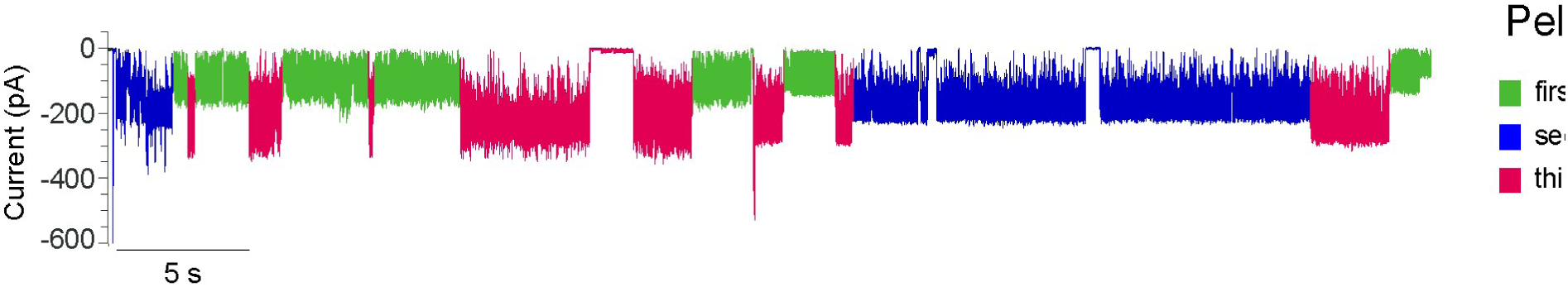
A representative ion current profile recorded in presence of multiple PelB molecules incorporated into the lipid bilayer.

**Supplemental Figure 15.**
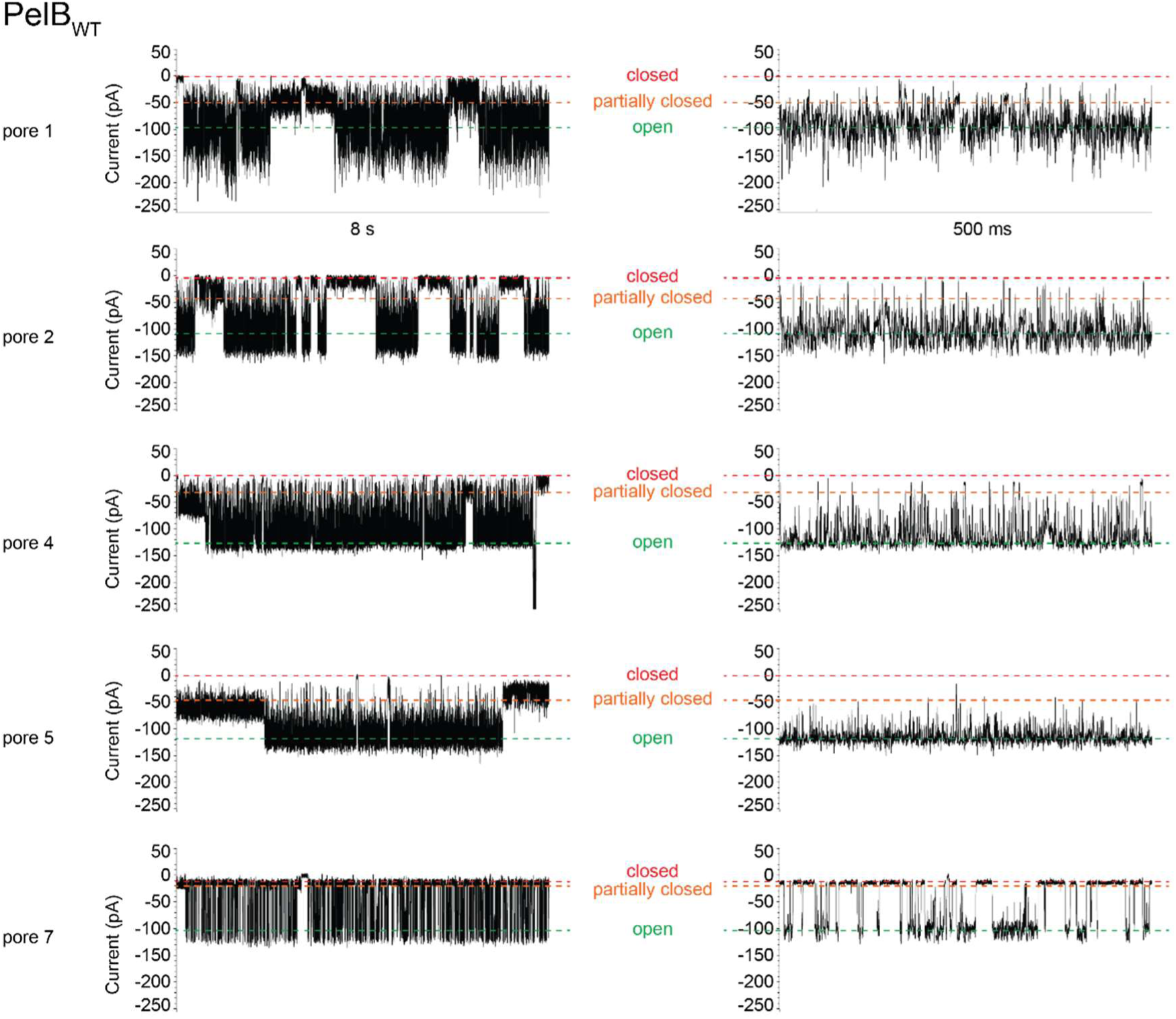
Single-channel recordings of wild-type PelB. Selected part of trace that depicted open pore current, as determined from the largest common denominating current of multiple pore insertions. Next to 8-s trace, a zoom-in of 500 ms is shown. A separate, partially closed/open state is also distinguished. Replicates share the open pore current size, but not open and closing frequency between the open, partially closed/open, and closed states. Experiments were conducted in 1M NaCl, 20 mM citric acid pH 3.4. Data was recorded at –150 mV applied potential, 50 kHz sampling rate and 10 kHz Bessel filter. Traces were additionally filtered with 2 kHz low-pass Gaussian filter.

**Supplemental Figure 16.**
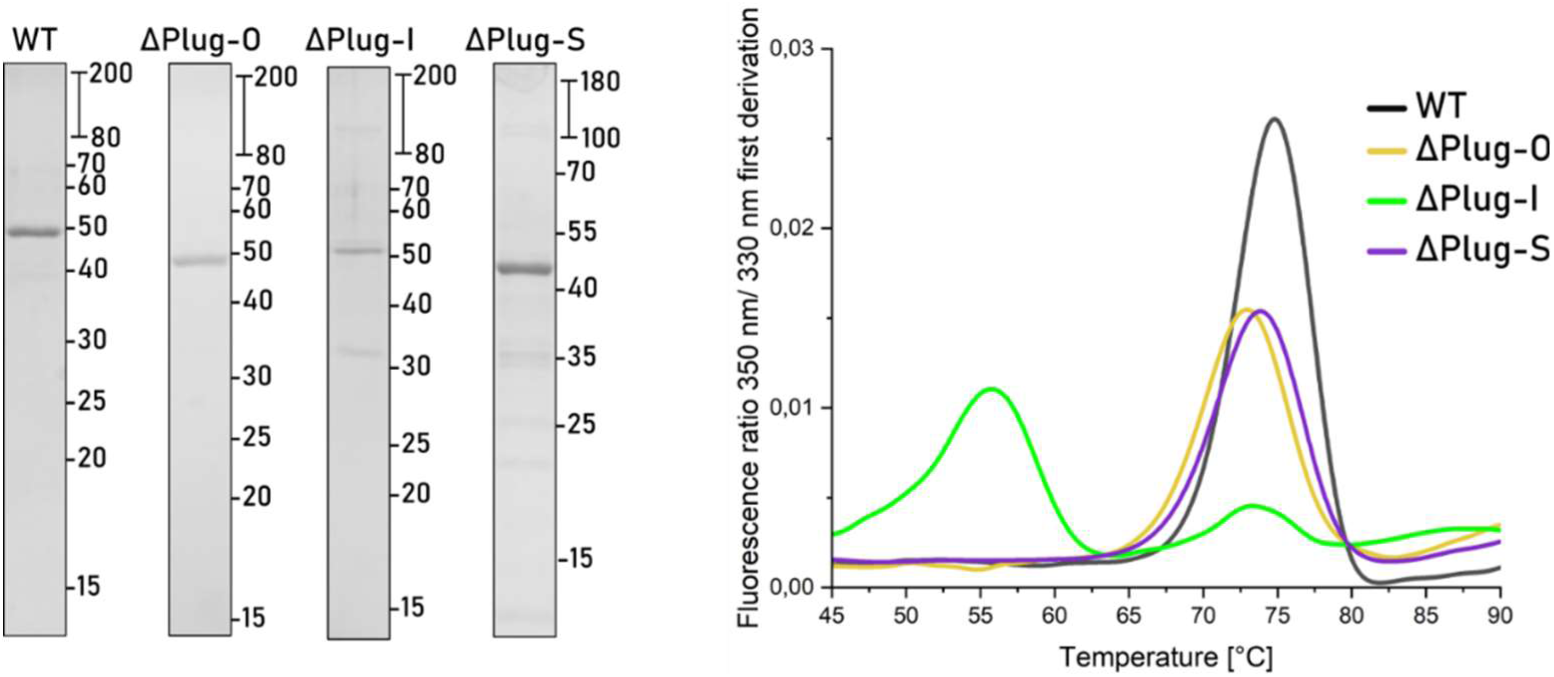
Left: Purification of PelB variants. Right: Thermal stability of the PelB variants in DDM tested by nanoDSF.

**Supplemental Figure 17.**
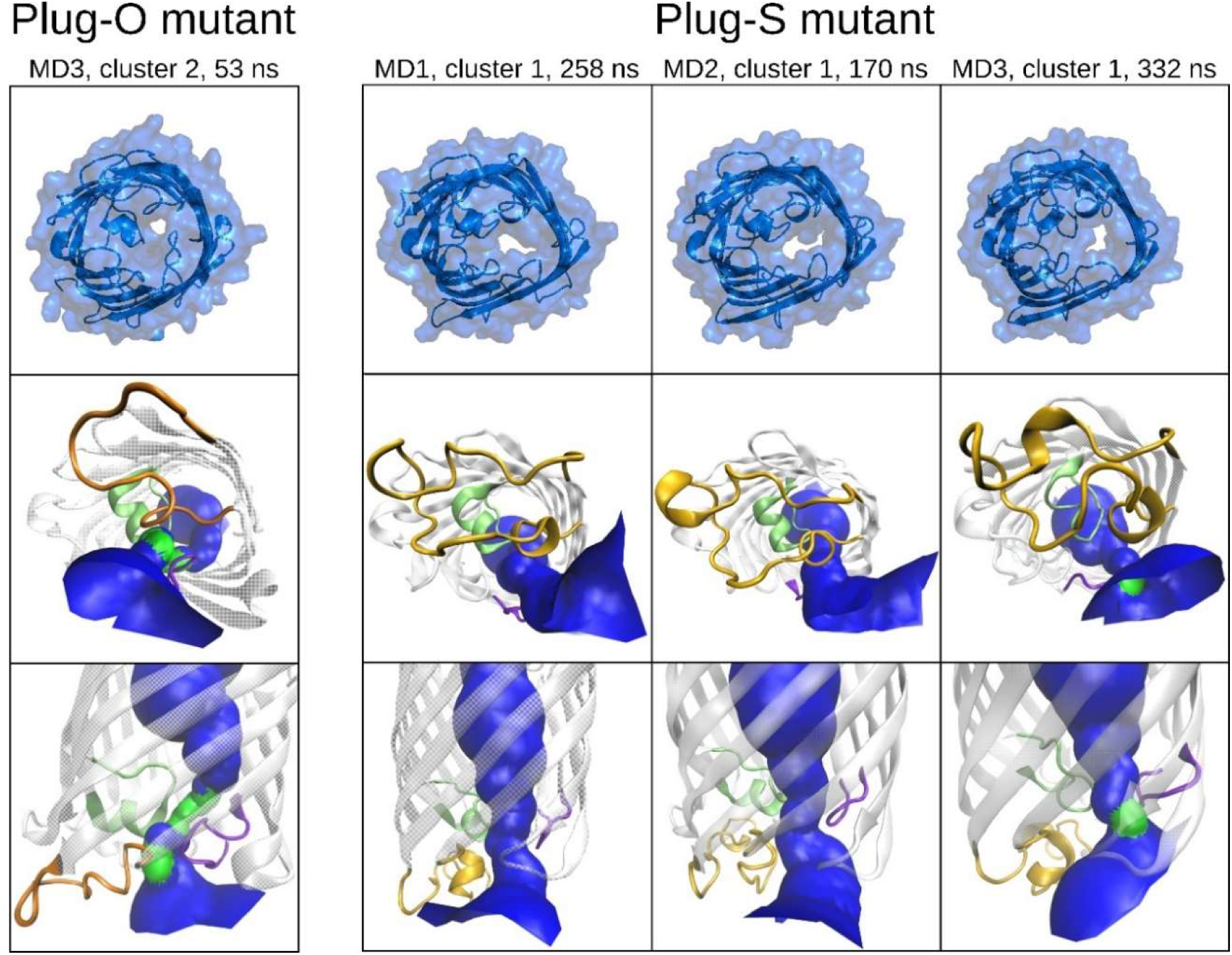
The pore analysis of PelB β-barrel the Plug-S and Plug-O mutant. The cartoon representation represents the orientation of the loops. The tunnel is colored in red for no water, in green for one water and blue for more water molecules, which are fitting in the pore. The pore analysis was done with HOLE algorithm and python package MDanalyis.

**Supplemental Figure 18:**
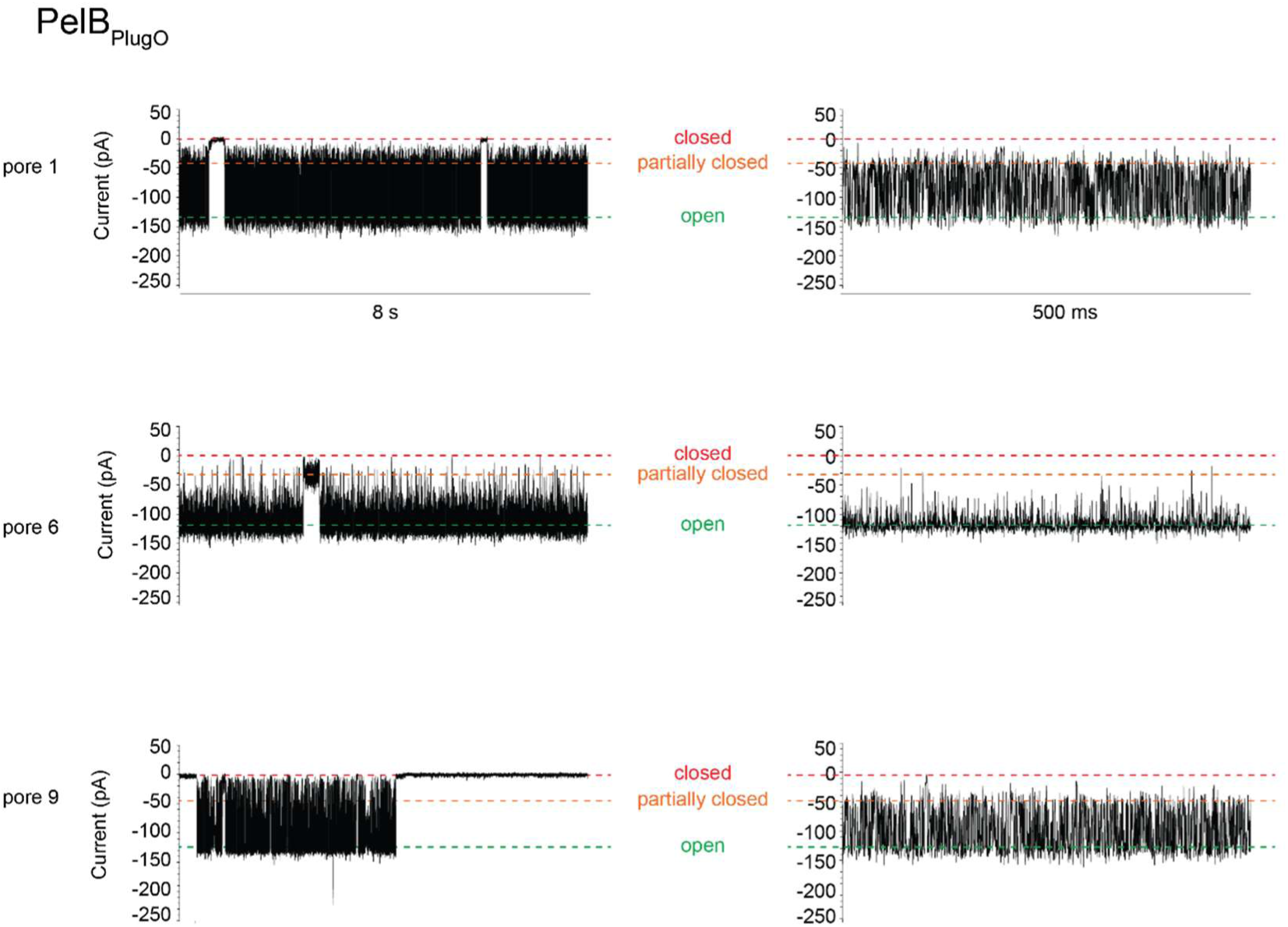
Single-channel recordings of PelB ΔPlug-O. Selected part of trace that depicted open pore current, as determined from the largest common denominating current of multiple pore insertions. Next to a 8 s trace, a zoom-in of 500 ms is shown. A separate, partially closed/open state is also distinguished. Replicates share the open pore current size, but not open and closing frequency between the open, partially closed/open, and closed states.

**Supplemental Figure 19.**
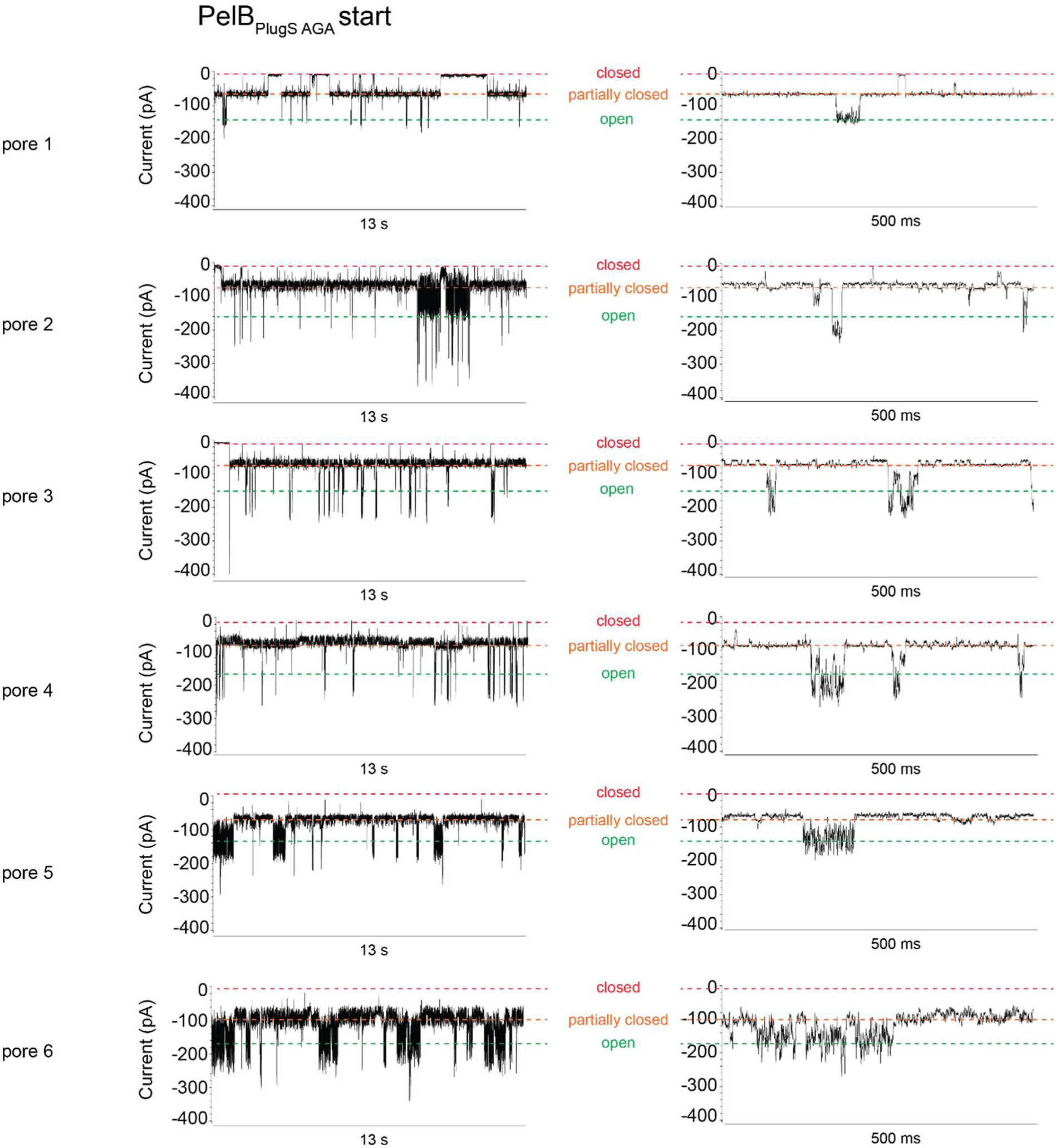
Single channel recordings of PelB ΔPlug-S. Representative parts of traces are shown. Next to a 13 s trace, a zoom-in of 500 ms is shown.

**Supplemental Figure 20.**
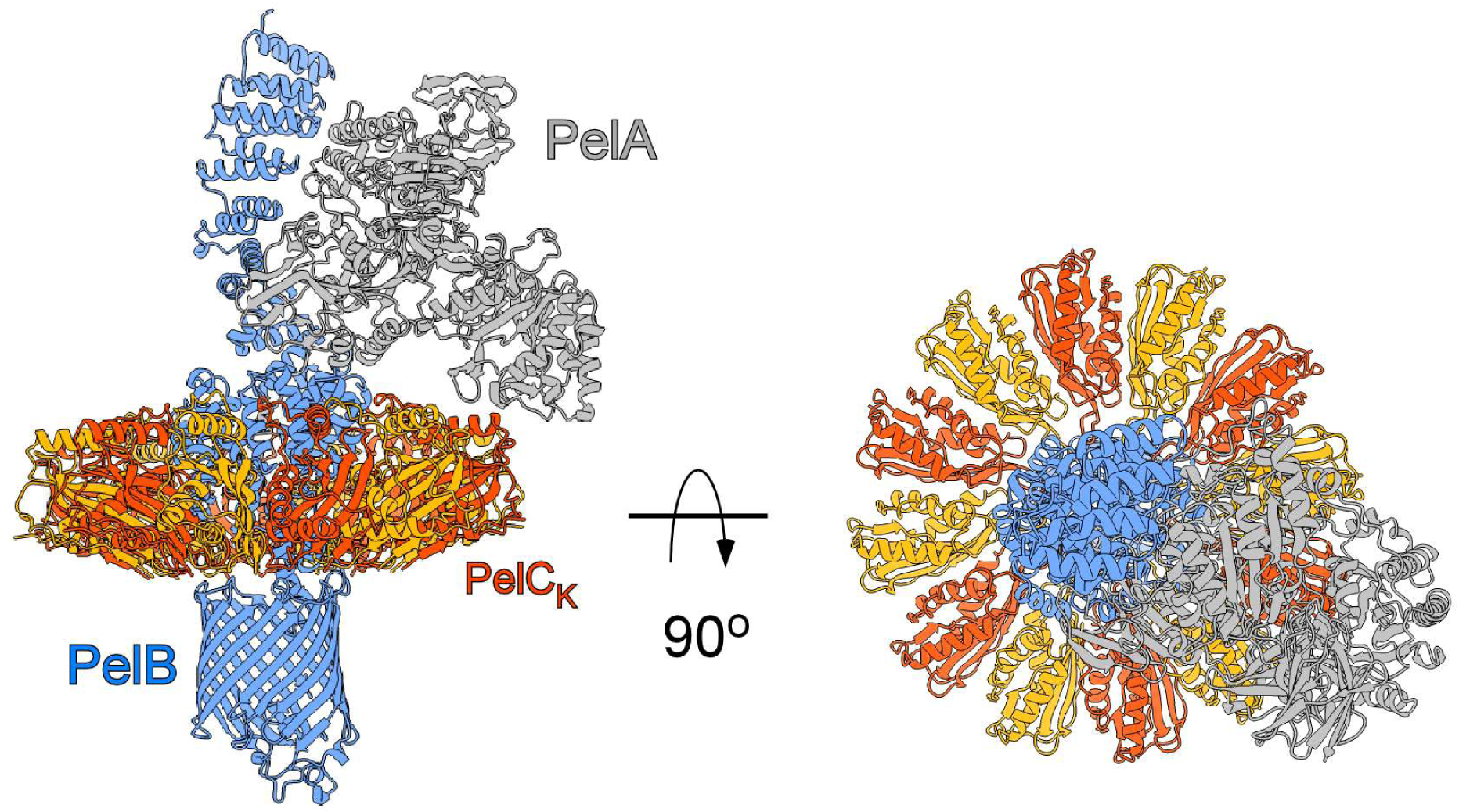
AlphaFold3-based model of the putative PelA-PelB-PelC complex. The hydrolase domain of PelB-bound PelA reaches PelC subunit K. PelB structure is shown from the residue 318.

## References

Abraham MJ, Murtola T, Schulz R, Páll S, Smith JC, Hess B, Lindahl E (2015) GROMACS: High performance molecular simulations through multi-level parallelism from laptops to supercomputers. SoftwareX 1–2: 19–25

Abramson J, Adler J, Dunger J, Evans R, Green T, Pritzel A, Ronneberger O, Willmore L, Ballard AJ, Bambrick J et al (2024) Accurate structure prediction of biomolecular interactions with AlphaFold 3. Nature 630: 493–500

Acheson JF, Derewenda ZS, Zimmer J (2019) Architecture of the Cellulose Synthase Outer Membrane Channel and Its Association with the Periplasmic TPR Domain. Structure 27: 1855–1861 e1853

Ansell TB, Song W, Coupland CE, Carrique L, Corey RA, Duncan AL, Cassidy CK, Geurts MMG, Rasmussen T, Ward AB et al (2023) LipIDens: simulation assisted interpretation of lipid densities in cryo-EM structures of membrane proteins. Nat Commun 14: 7774

Daura X, Gademann K, Jaun B, Seebach D, Van Gunsteren WF, Mark AE (1999) Peptide Folding: When Simulation Meets Experiment. Angewandte Chemie International Edition 38: 236–240

Gheorghita AA, Wozniak DJ, Parsek MR, Howell PL (2023) Pseudomonas aeruginosa biofilm exopolysaccharides: assembly, function, and degradation. FEMS Microbiol Rev 47

Goyal P, Krasteva PV, Van Gerven N, Gubellini F, Van den Broeck I, Troupiotis-Tsailaki A, Jonckheere W, Pehau-Arnaudet G, Pinkner JS, Chapman MR et al (2014) Structural and mechanistic insights into the bacterial amyloid secretion channel CsgG. Nature 516: 250–253

Hagn F, Etzkorn M, Raschle T, Wagner G (2013) Optimized phospholipid bilayer nanodiscs facilitate high-resolution structure determination of membrane proteins. J Am Chem Soc 135: 1919–1925

Huang J, MacKerell AD, Jr. (2013) CHARMM36 all-atom additive protein force field: validation based on comparison to NMR data. J Comput Chem 34: 2135–2145

Huang J, Rauscher S, Nawrocki G, Ran T, Feig M, de Groot BL, Grubmuller H, MacKerell AD, Jr. (2017) CHARMM36m: an improved force field for folded and intrinsically disordered proteins. Nat Methods 14: 71–73

Jennings LK, Storek KM, Ledvina HE, Coulon C, Marmont LS, Sadovskaya I, Secor PR, Tseng BS, Scian M, Filloux A et al (2015) Pel is a cationic exopolysaccharide that cross-links extracellular DNA in the Pseudomonas aeruginosa biofilm matrix. Proc Natl Acad Sci U S A 112: 11353–11358

Jo S, Kim T, Iyer VG, Im W (2008) CHARMM-GUI: a web-based graphical user interface for CHARMM. J Comput Chem 29: 1859–1865

Josts I, Stubenrauch CJ, Vadlamani G, Mosbahi K, Walker D, Lithgow T, Grinter R (2017) The Structure of a Conserved Domain of TamB Reveals a Hydrophobic beta Taco Fold. Structure 25: 1898–1906 e1895

Kanonenberg K, Royes J, Kedrov A, Poschmann G, Angius F, Solgadi A, Spitz O, Kleinschrodt D, Stuhler K, Miroux B et al (2019) Shaping the lipid composition of bacterial membranes for membrane protein production. Microb Cell Fact 18: 131

Le Mauff F, Razvi E, Reichhardt C, Sivarajah P, Parsek MR, Howell PL, Sheppard DC (2022) The Pel polysaccharide is predominantly composed of a dimeric repeat of alpha-1,4 linked galactosamine and N-acetylgalactosamine. Commun Biol 5: 502

Lee J, Patel DS, Stahle J, Park SJ, Kern NR, Kim S, Lee J, Cheng X, Valvano MA, Holst O et al (2019) CHARMM-GUI Membrane Builder for Complex Biological Membrane Simulations with Glycolipids and Lipoglycans. J Chem Theory Comput 15: 775–786

Li A, Schertzer JW, Yong X (2018) Molecular dynamics modeling of Pseudomonas aeruginosa outer membranes. Phys Chem Chem Phys 20: 23635–23648

Liang B, Tamm LK (2007) Structure of outer membrane protein G by solution NMR spectroscopy. Proc Natl Acad Sci U S A 104: 16140–16145

Lindahl A, Hess, & van der Spoel, GROMACS 2021.6 Source code (2021.5). Zenodo.

Marmont LS, Rich JD, Whitney JC, Whitfield GB, Almblad H, Robinson H, Parsek MR, Harrison JJ, Howell PL (2017a) Oligomeric lipoprotein PelC guides Pel polysaccharide export across the outer membrane of Pseudomonas aeruginosa. Proc Natl Acad Sci U S A 114: 2892–2897

Marmont LS, Whitfield GB, Rich JD, Yip P, Giesbrecht LB, Stremick CA, Whitney JC, Parsek MR, Harrison JJ, Howell PL (2017b) PelA and PelB proteins form a modification and secretion complex essential for Pel polysaccharide-dependent biofilm formation in Pseudomonas aeruginosa. J Biol Chem 292: 19411–19422

Mirdita M, Schutze K, Moriwaki Y, Heo L, Ovchinnikov S, Steinegger M (2022) ColabFold: making protein folding accessible to all. Nat Methods 19: 679–682

Morgan JL, Strumillo J, Zimmer J (2013) Crystallographic snapshot of cellulose synthesis and membrane translocation. Nature 493: 181–186

Nelson RE, Hatfield KM, Wolford H, Samore MH, Scott RD, Reddy SC, Olubajo B, Paul P, Jernigan JA, Baggs J (2021) National Estimates of Healthcare Costs Associated With Multidrug-Resistant Bacterial Infections Among Hospitalized Patients in the United States. Clin Infect Dis 72: S17–S26

Nikaido H, Rosenberg EY (1983) Porin channels in Escherichia coli: studies with liposomes reconstituted from purified proteins. J Bacteriol 153: 241–252

Punjani A, Rubinstein JL, Fleet DJ, Brubaker MA (2017) cryoSPARC: algorithms for rapid unsupervised cryo-EM structure determination. Nat Methods 14: 290–296

Ritchie TK, Grinkova YV, Bayburt TH, Denisov IG, Zolnerciks JK, Atkins WM, Sligar SG (2009) Chapter 11 - Reconstitution of membrane proteins in phospholipid bilayer nanodiscs. Methods Enzymol 464: 211–231

Suits MD, Sperandeo P, Deho G, Polissi A, Jia Z (2008) Novel structure of the conserved gram-negative lipopolysaccharide transport protein A and mutagenesis analysis. J Mol Biol 380: 476–488

Wang Y, Andole Pannuri A, Ni D, Zhou H, Cao X, Lu X, Romeo T, Huang Y (2016) Structural Basis for Translocation of a Biofilm-supporting Exopolysaccharide across the Bacterial Outer Membrane. J Biol Chem 291: 10046–10057

Whitfield GB, Marmont LS, Bundalovic-Torma C, Razvi E, Roach EJ, Khursigara CM, Parkinson J, Howell PL (2020a) Discovery and characterization of a Gram-positive Pel polysaccharide biosynthetic gene cluster. PLoS Pathog 16: e1008281

Whitfield GB, Marmont LS, Ostaszewski A, Rich JD, Whitney JC, Parsek MR, Harrison JJ, Howell PL (2020b) Pel Polysaccharide Biosynthesis Requires an Inner Membrane Complex Comprised of PelD, PelE, PelF, and PelG. J Bacteriol 202

Whitney JC, Hay ID, Li C, Eckford PD, Robinson H, Amaya MF, Wood LF, Ohman DE, Bear CE, Rehm BH et al (2011) Structural basis for alginate secretion across the bacterial outer membrane. Proc Natl Acad Sci U S A 108: 13083–13088

Willems K, Van Meervelt V, Wloka C, Maglia G (2017) Single-molecule nanopore enzymology. Philos Trans R Soc Lond B Biol Sci 372

Wu EL, Cheng X, Jo S, Rui H, Song KC, Davila-Contreras EM, Qi Y, Lee J, Monje-Galvan V, Venable RM et al (2014) CHARMM-GUI Membrane Builder toward realistic biological membrane simulations. J Comput Chem 35: 1997–2004

Zheng SQ, Palovcak E, Armache JP, Verba KA, Cheng Y, Agard DA (2017) MotionCor2: anisotropic correction of beam-induced motion for improved cryo-electron microscopy. Nat Methods 14: 331–332

